# Photocontrol of endogenous glycine receptors *in vivo*

**DOI:** 10.1101/744391

**Authors:** Alexandre M.J. Gomila, Karin Rustler, Galyna Maleeva, Alba Nin-Hill, Daniel Wutz, Antoni Bautista-Barrufet, Xavier Rovira, Miquel Bosch, Elvira Mukhametova, Marat Mukhamedyarov, Frank Peiretti, Mercedes Alfonso-Prieto, Carme Rovira, Burkhard König, Piotr Bregestovski, Pau Gorostiza

## Abstract

Glycine receptors (GlyRs) are indispensable to maintain excitatory/inhibitory balance in neuronal circuits controlling reflex and rhythmic motor behaviors. Here we have developed Glyght, the first GlyR ligand controlled with light. It is selective over other cys-loop receptors, active *in vivo*, and displays an allosteric mechanism of action. The photomanipulation of glycinergic neurotransmission opens new avenues to understand inhibitory circuits in intact animals, and to develop drug-based phototherapies.

## MAIN

The control of biological activity with light has become a powerful tool to understand complex multicellular^1^ and intracellular processes^2^, protein dynamics^3, 4^, as well as to develop novel therapeutic strategies^4^. In particular, optogenetics^5^ and photopharmacology^6^ have been a boon for neurobiology, empowering it to control neuronal receptor activity with subtype pharmacological selectivity and micrometric resolution^7, 8^, firing individual neurons^9^ and mapping their connectivity and strength^10^. Photoswitchable ligands^6^ enable directly controlling the activity of endogenous receptors without requiring genetic manipulation^8, 11^. They can be applied to intact tissue, making drug-based phototherapies possible. Despite the importance of inhibitory receptors, few photoswitches targeting ionotropic gamma amino butyric acid receptors (GABA_A_Rs) have been reported^12–15^, and no specific modulator has been developed for glycine receptors (GlyRs).

GlyRs and GABA_A_Rs belong to the pentameric Cys-loop superfamily along with excitatory nicotinic acetylcholine (nAChRs) and serotonin receptors (5-HT_3_R). GABA_A_Rs and GlyRs share not only the pentameric assembly of their subunits and the inhibitory regulation of cell membrane potential through their chloride-selective pore, but also the mechanism of agonist-induced desensitization, which is driven by homologous residues between transmembrane domains of the receptor^16^. These similarities hamper the development of selective ligands.

Alterations in inhibitory neurotransmission cause an excitation/inhibition disbalance that has been linked to many and etiologically diverse neurological diseases, from epilepsy to anxiety, autism^17^ and schizophrenia^18^. Modulating neuronal inhibition with systemically administered drugs can partially restore balance and reduce certain symptoms, but the efficacy of these strategies is very limited in diseases where specific circuits are altered. In these cases, what matters is the precise location and timing of inhibition to restore homeostasis, and cure is unlikely even with the most pharmacologically selective drugs.

In particular, glycinergic transmission regulates the majority of reflex and rhythmic motor behaviors, including locomotion and breathing^19^. Normal functioning of locomotor circuits relies on a strictly determined equilibrium between excitatory and inhibitory synapses onto interneurons^20^. Disruptions to this balance trigger locomotor dysfunctions^21^. Insufficient GlyR function leads to excessive startle response (hyperekplexia) and other pathologies^22, 23^. Despite the importance of glycinergic neurotransmission, the repertoire of GlyR drugs available is still extremely limited^24, 25^ and not highly specific^26^. Strychnine and tropisetron remain the only modulators that show selectivity for GlyR over GABA_A_R, but the former is also nAChR antagonist^27, 28^, and the latter antagonizes 5-HT_3_R^29, 30^. Selective GlyR drugs are necessary to treat hyperekplexia, autism, chronic inflammatory pain, breathing disorders, temporal lobe epilepsy, alcoholism, and motor neuron disease^26^. However, given the diversity and ubiquity of glycinergic circuits, traditional pharmacology is unlikely to be enough unless the activity of these ligands can be modulated right at the specific circuits involved in every disease.

Here, we initially aimed at developing photoswitchable ligands of GABA_A_Rs based on benzodiazepines. Azo-compounds (**3a-d**, **Figure 1a**) were thus designed to display *cis-trans* photochromism while maintaining the ability of the nitrazepam moiety to bind the GABA_A_R, as reported for other substitutions at the same position of the ligand^31^. Unexpectedly, we obtained a GlyR-selective negative allosteric modulator whose inhibitory action at GlyRs was increased under UV light (*cis*-on), and that we named Glyght (short for ‘<u>Gly</u>R controlled by li<u>ght</u>’).

**Fig. 1:**
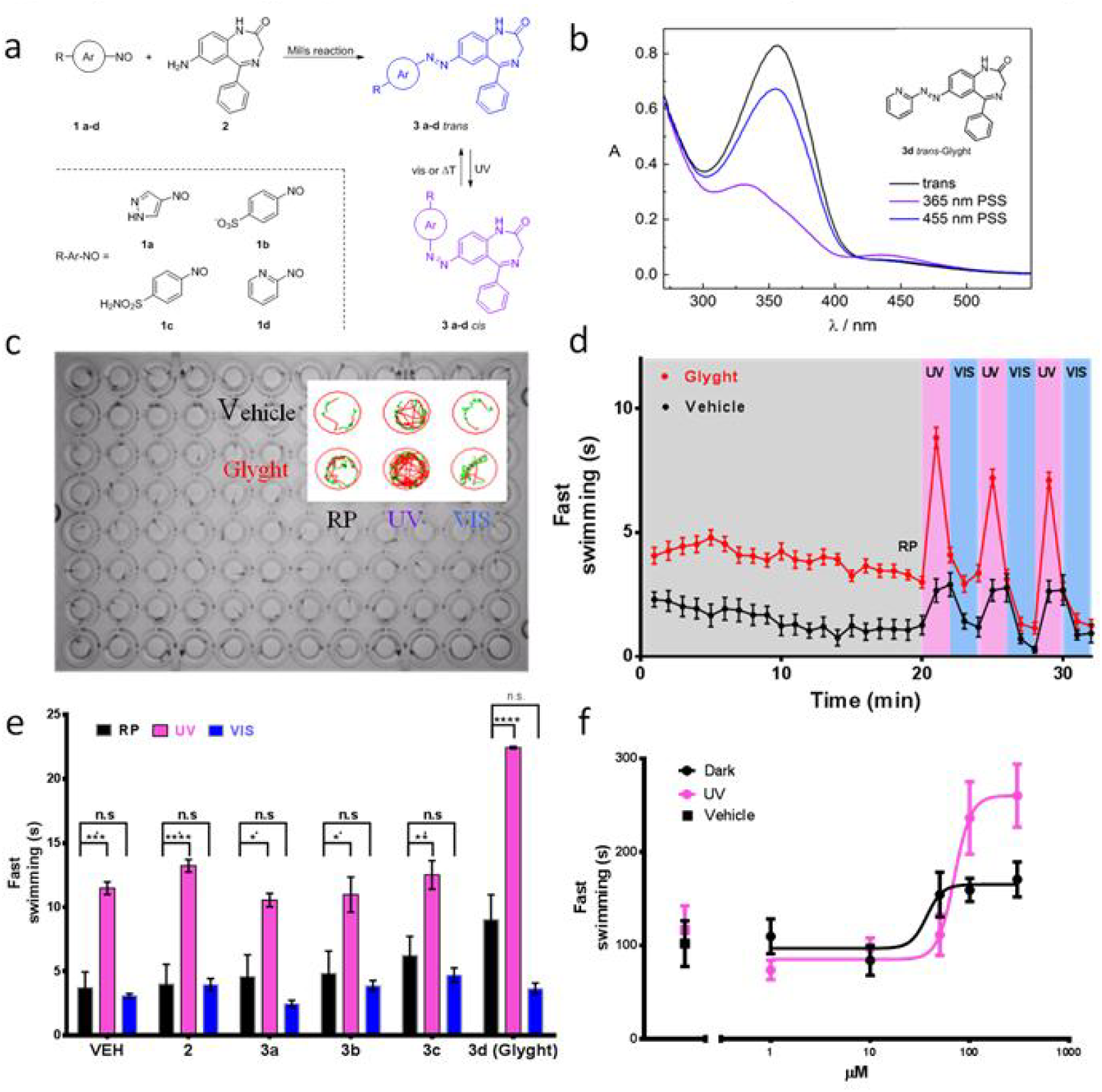
Synthetic strategy of photochromic derivatives of benzodiazepine and behavioral assay. **a,** General scheme for synthesizing azobenzene derivatives of benzodiazepine with different substitutions via Mills reaction (3a-d). The resulting trans-cis photoisomerization is indicated. **b,** UV-visible absorption spectrum showing the photochromic behavior of compound **3d (Glyght)**. **c,** 96-well plate with individual zebrafish larvae in each well exposed to different solutions conditions. Video recordings of the entire plate under different illumination conditions (dark, 365 nm, 455 nm) allow analyzing the motility of individual animals in order to identify drug- and light-dependent activities. Inset: Detail of two wells containing vehicle (1% DMSO, top) and compound **3d** (**Glyght**, bottom) and the trajectories swam by individual larvae during the resting, ultraviolet, and visible periods (RP, UV, VIS, respectively). Different swimming velocities during each trajectory are automatically categorized by the software and are indicated in green (2-6 mm·s^-1^) and red (faster than 6 mm·s^-1^). **d,** Fast swimming (FS, time spent swimming faster than 6 mm·s^-1^) of individual larvae exposed to vehicle and Glyght during RP, UV and VIS illumination was integrated into 1 min data points. Error bars indicate the standard deviation of the mean (S.E.M.) of FS in each 1 min period for 12 individual larvae. **e,** Quantification of larvae activity (n=8) during RP (averaged over 2 min) and during UV and visible periods (averaged over 3 cycles) for 7-nitrazepam (7NA (**2**), non-photoswitchable GABA_A_R inhibitor), vehicle and a panel of 4 photochromic ligands. Glyght displays significantly higher activity during RP and UV periods, suggesting basal and UV-enhanced antagonism of inhibitory receptors. Error bars represent S.E.M. **f,** Dose response for Glyght in FS (in seconds) according to the last 2 minutes of RP (Dark trace) and 2 minutes of 365 nm illumination (UV trace). Dark trace activity corresponds to pure *trans*-Glyght and UV trace to isomerized *cis*-Glyght. Error bars represent S.E.M. for n = 8 larvae per concentration group and traces fitted with sigmoidal four parameters model.

The photochromic azo moiety was introduced via a Mills reaction of several nitroso aryl moieties (compounds **1a-d**, **Figure 1a**) with the amino-substituted benzene in the structure of 7-aminonitrazepam (compound **2**, **Figure 1a**, obtained as reported^32, 33^). All derivatives reversibly photoswitch at 365 nm (*trans* to *cis* isomerization) and 455 to 530 nm (*cis* to *trans*isomerization), as exemplarily shown for the pyridine-based derivative **3d** (Glyght) (**Figure 1b**). Thermal relaxation half-lives in the dark are longer than 1h (**Supplementary Table 1** and **Supplementary Figures 19-22**).

To characterize the photopharmacological effects of compounds **3a-d**, we designed a behavioral assay to record and quantify the swimming activity of zebrafish larvae as a function of illumination. Zebrafish express all GABA_A_R and GlyR subunits, with high sequence similarity to the mammalian receptors^34^, and larvae display full exploratory capacities at 7 days post fertilization (dpf)^35–37^. Ligands of inhibitory receptors should alter the well known behavioural activity of the larvae, and be correlated to specific dynamic traits such as speed swimming variations, transition swimming patterns or anxiety-like behaviours^38, 39^. In order to identify alterations in inhibitory neurotransmission, we focused on fast movements and measured swimming distances and duration of high speed swimming^38, 40^. Individual larvae were placed in separate wells of a 96-well plate (**Figure 1c**), each containing different solution conditions including non-photoswitchable control drugs like GABA_A_R potentiator 7-amino nitrazepam (**2**, 7AN), the photochromic compounds **3a-d** at different concentrations, and vehicle (DMSO 1%). The setup (see online Methods for details) allows maintaining the animals in the dark and subjecting them to cycles of illumination at 365 and 455 nm. The inset of **Figure 1c** shows exemplarily 1-minute trajectories of individual fish in wells containing vehicle and compound **3d** (Glyght, 100μM) during the resting period (RP), under 365 nm (ultraviolet, UV) and under 455 nm (visible, VIS). Green and red trajectories plot slow and fast swimming periods, respectively. Videos of the entire plate during the RP and photoswitching experiment can be viewed in the Supplementary Movie 1. The time course (integrated every minute for 12 animals) is shown in **Figure 1d**. During the RP, Glyght-treated animals display enhanced locomotion compared to controls. Although control animals are startled by UV light and slowed down by visible light, the increase in locomotion displayed by treated animals is significantly higher. The results of all photoswitches (**3a-d**), 7AN (**2**), and vehicle are shown in **Figure 1e**. In all cases, the time spent in fast swimming (see online Methods for details) is longer under UV than under visible light, but only for Glyght (**3d**) this difference is higher than for controls. In order to identify photoswitchable hits, we assumed larvae swimming activity as the quantifiable variable, and we defined light periods as intrinsically dependent variables of larvae behavioral outcomes. We calculated the ratio between the activities under UV and under visible light (UV/vis activity ratio, UVAR) and used it as a score to identify compounds producing photoswitchable behaviors. Three compounds displayed UVARs significantly different from the control (UVAR of endogenous photoresponses in vehicle, **Supplementary Figure 1**). One was excluded due to precipitation over time (**3c**) and two were retained for further studies: **3b** (azo-NZ1, a GABA_A_R blocker reported in a separate article^41^), and **3d** (Glyght) characterized below. For Glyght, we confirmed the hit in 5 independent experiments from different larvae batches (**Supplementary Figures 2** and **3**) and verified that photoresponses are dosedependent (**Figure 1e**) before moving on to *in vitro* pharmacological characterizations with an activity assay.

We used electrophysiological recordings to measure agonist-induced responses in anion-selective GABA_A_Rs and GlyRs, and in cation-selective 5-HT_3_Rs and AChRs (see online Methods for details), and evaluated the effects of adding Glyght. The results are shown in **Figure 2abc** and **Supplementary Figures 4-8**. In contrast to the effect of diazepam in GABA_A_Rs^42^, Glyght displayed a weak action on α_1_/β_2_/γ_2_ GABA_A_Rs: in the presence of 50 μM *trans*-Glyght, 5μM GABA-induced currents were inhibited only by 9 ± 2% (n = 6) and 300μM GABA currents by 14 ± 4% (n = 5) (**Supplementary Figure 4**). UV light prevented the weak current inhibition of GABA_A_R by Glyght.

**Fig. 2:**
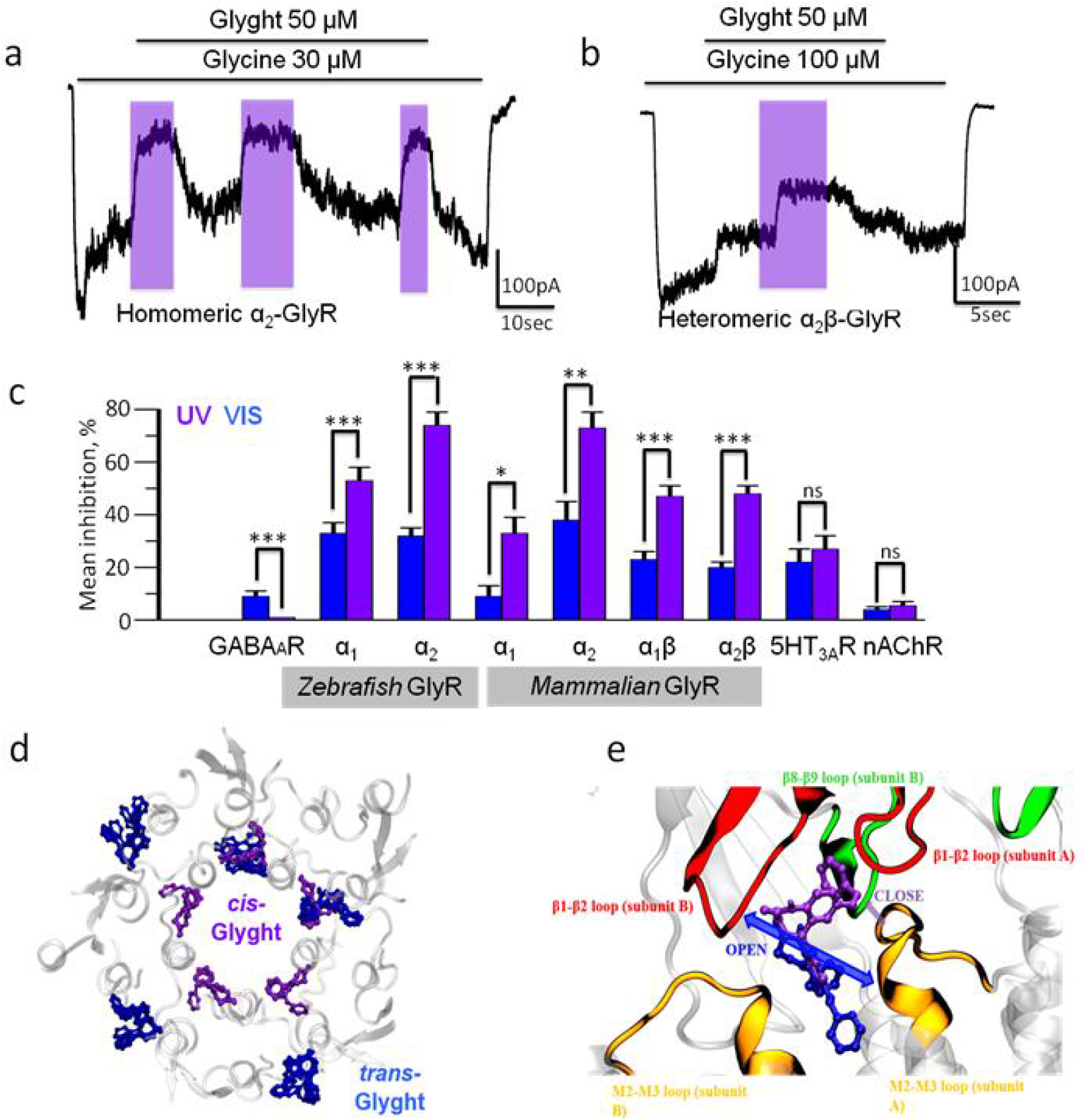
Glyght is a GlyR selective photoswitch. **a,** Representative trace illustrating the photoswitchable effect of Glyght (50 μM) on currents mediated by GlyR composed of α_2_ subunits. Note an increase of the inhibitory efficiency of Glyght upon illumination with UV light. The duration of glycine and Glyght application is indicated by black bars above traces; the duration of UV illumination is indicated by violet rectangles (V_hold_ = −30 mV). **b,** Representative trace illustrating the photoswitching action of Glyght (50 μM) on currents mediated by heteromeric α_2_/βGlyRs (V_hold_ = −30mV). **c,** Cumulative graph demonstrating the percentage of Glyght (50μM) induced inhibition under visible (blue column) and UV (violet) light on different types of Cys-loop receptors. *- p ≤ 0.05, **- p ≤ 0.01, ***- p ≤ 0.001. **d,** Molecular modeling results, where the five α_1Z_GlyR subunits are colored in white and represented in ribbons, and the most populated poses obtained in the flexible docking of *cis-* (violet) and *trans*-Glyght (blue) in α_1Z_GlyR are represented in ball and sticks. Glyght binds preferentially at a non-glycine site located at the interface of the extracellular and transmembrane domains (ECD and TMD) that is involved in the allosteric coupling between ligand binding to the ECD and opening of the ion channel pore in the TMD^44-48^. **e,** Detailed view of the intersubunit site at the ECD/TMD interface including the protein loops changing conformation upon receptor activation^49^. Interactions of the M2-M3 loop with the β8-β9 and β1-β2 loops are associated with stabilization of the closed (purple arrow) and open (blue arrow) states, respectively^49^. *Cis*-Glyght links the M2-M3 and β8-β9 loops, thus favouring the closed state (purple arrow). These results are in full agreement with stronger inhibition of GlyRs observed for *cis*-Glyght in panels **abc**. In contrast, *trans-Glyght* may interfere with the interaction between the M2-M3 and β1-β2 loops (blue arrow) and thus with open state stabilization.

Since the action of Glyght on GABA_A_Rs cannot account for the robust behavioral effects observed in fish (**Figure 1c-f**), we asked whether this compound might interact with other inhibitory ligand-gated receptors responsible for the control of movement^26, 43^, namely GlyR. Thus, we tested Glyght in zebrafish α_1_ (α_1Z_) and α_2_ (α_2Z_) homomeric GlyRs during activation by a non-saturating concentration of glycine (see online Methods). In α_1Z_GlyRs, *trans*-Glyght (50 μM) reduced the current amplitude by 33 ± 4% (**Supplementary Figure 5a**) and stronger reduction was observed upon isomerizing it to *cis*-Glyght under 365 nm light (53 ± 5%, n = 8). Receptor photoswitching was stronger on α_2Z_GlyRs wherein 50 μM *trans*-Glyght and *cis*-Glyght reduced glycine currents by 32 ± 3% and 74 ± 5% respectively (**Figure 2c** and **Supplementary Figure 5def**, n = 7). When Glyght was co-applied with saturating concentrations of glycine, its effect become negligible, clearly indicating that the compound is not an open channel blocker (**Supplementary Figure 5bc**, n = 3; **ef**, n = 3). Full dose-response curves under UV and dark conditions in these receptors confirmed that α_2Z_GlyRs are more sensitive to Glyght than α_1Z_GlyRs (**Supplementary Figure 5hi**). Importantly, these results were confirmed in mammalian GlyRs where Glyght caused strong *cis*-on inhibition in all homomeric and heteromeric receptors (α_1_, α_2_, α_1_β, and α_2_β; **Figure 2ab** and **Supplementary Figure 5abde**). No illumination-dependent outcome was observed using other inhibitors like picrotoxin (**Supplementary Figure 6cf**). Since Glyght displays selectivity for GlyRs compared to GABA_A_Rs (**Figure 2c**), we further characterized its activity in other pentameric receptors. The compound displayed low activity and no photoswitching in 5-HT_3A_R (**Figure 2c** and **Supplementary Figure 7**) and was completely inactive in muscular nAChR (**Supplementary Figure 8**). The results are summarized in **Figure 2c** and indicate that Glyght is broadly active in homo- and heteromeric GlyRs from fish and mammals, and remarkably selective *versus* all other members of the Cys-loop receptor family.

To understand the photopharmacological profile of Glyght we turned to modeling of the compound in the open (agonist-bound) structure of α_1Z_GlyR^49^. Molecular dockings showed that neither of the Glyght isomers can bind at the channel pore (in agreement with our patch clamp results), that *trans*-Glyght displays moderate binding in several regions of the extracellular and transmembrane domains (ECD and TMD), and that *cis*-Glyght poses to a non-glycine site at the ECD/TMD interface (displayed in blue and violet respectively, **Figure 2de** and **Supplementary Figure 9-10**). We focused on the latter binding site, since it is the region showing the largest differences in ligand pose densities for both α1_Z_ and α2_H_GlyRs. Moreover, this site includes key residues for channel activation and conductance^50, 51^ and is involved in allosteric coupling between ligand binding to the ECD and ion channel pore opening in the TMD^44-48^. As shown in **Figure 2e**, *cis*-Glyght binds further inside the ECD/TMD interface than *trans*-Glyght, in line with its stronger effect. Moreover, *trans*-Glyght can mediate the interaction between M2-M3 and β1-β2 loops that stabilizes the open channel state^49^ (blue arrow in **Figure 2e**). On the other hand, *cis*-Glyght favors the interaction between M2-M3 and β8-β9 loops, which are associated to the closed state (purple arrow in **Figure 2e**). These results are in full agreement with the stronger inhibition of GlyRs observed for *cis*-Glyght in **Figure 2abc**.

The optogenetic control of excitatory and inhibitory circuits is defined genetically based on available cell-specific promoters^52^. Photoswitchable receptor-selective drugs are powerful complementary tools that enable spatiotemporal control over pharmacologically defined circuits (glutamatergic, cholinergic, gabaergic, glycinergic), and cast new light on classic systems neuroscience research and drug-based therapies. We have used a zebrafish behavioral assay and photoresponse score to identify the first GlyR-selective, light-regulated inhibitor. Glyght loses inhibitory activity at saturating glycine concentrations, which is compatible with orthosteric or allosteric antagonism^53^ (**Supplementary Figure 5bcef**). Molecular modeling results indicate that Glyght binds to an allosteric site that regulates GlyR gating and conductance^44–48, 50, 51^ (**Figure 2de**). Thus, Glyght also provides a template to design new allosteric non-photoswitchable ligands like amide, ether or methoxy analogs bearing general pharmacological interest. Glyght reversibly elicits excitatory behaviors in zebrafish larvae and is a good candidate to study glycinergic neurotransmission in spatiotemporally defined patterns, and to explore therapeutic approaches based on localized and selective activation of GlyRs. The partial activity of Glyght in 5-HT_3A_Rs is consistent with tropisetron’s effect in both receptors^30^ and anticipates moderate emetic activity. However, its high selectivity with respect to nicotinic receptors makes Glyght remarkably superior to strychnine^27^, raising hopes to modulate GlyRs without concomitant toxicity and opening a new avenue to clinical pharmacology at large.

## Acknowledgements

We are grateful to S. Lummis for the 5□HT3A subunit cDNA. This study was supported by ERA SynBIO Grant MODULIGHTOR (PCIN□2015□163□C02□01), the Russian Science Foundation (grant 18□15□00313), AGAUR/Generalitat de Catalunya (CERCA Programme 2017□SGR□1442), FEDER funds, Human Brain Project WAVESCALES (SGA2 Grant Agreement 785907), Fundaluce Foundation, MINECO (Project CTQ2016□80066R), a FPI-MICIU Ph.D. scholarship to A. M. J. G., a FI□AGAUR Ph.D. scholarship to A. N.□H., and an IBEC-BEST postdoctoral scholarship to G. M. We also thankfully acknowledge the computer resources at MareNostrum III and MinoTauro and the technical support provided by the Barcelona Supercomputing Center (BCV-2017-2-0004).

## Author contributions

A.M.J.G and X.R. performed *in vivo* experiments. K.R., D.W. and A.B.B. performed compounds chemical synthesis and characterisation. G.M., E.M., M.M, F.K. and M.B. performed *in vitro* experiments. A.N.H and M.P. performed molecular modeling simulations and analysis. C.R. supervised molecular modeling. B.K. supervised chemical synthesis. P.B. supervised *in vitro* experiments. P.G. conceived the project and supervised *in vivo* experiments. AMJG and PG wrote the manuscript with contributions from all authors.

## Competing interests

The authors declare no competing interests.

