## Supplementary Information for "Photocontrol of endogenous glycine receptors *in vivo*"

### METHODS

#### Chemistry and photochromism

7-aminonitrazepam (**2**)[1,2] and the nitroso derivatives **1a**[3], **1b**[4], **1c**[5], and **1d**[6]were synthesized according to reported procedures. Commercial reagents and starting materials were purchased from Acros Organics, Alfa-Aesar, Fisher Scientific, Sigma Aldrich or VWR and used without any further purification. Solvents were used in p.a. quality and dried according to common procedures, if necessary. Commercially phosphate buffer (pH = 7.4) was used for investigations of the photochromic properties. Dry nitrogen was used as inert gas atmosphere. A Biotage Isolera flash purification system with UV/Vis detector using Sigma Aldrich MN silica gel 60 M (40-63 µm, 230-400 mesh) for normal phase or pre-packed Biotage SNAP cartridges (KP C18 HS) for reversed phase chromatography was used for automated flash column chromatography. Reaction monitoring via TLC and determination of $R_f$ values was accomplished on alumina plates coated with silica gel (Merck silica gel 60 F254, 0.2 mm). Melting points were measured with a Stanford Research Systems OptiMelt MPA 100 device and are uncorrected. NMR spectra were measured on BrukerAvance 300 ($^1$H 300.13 MHz, $^{13}$C 75.48 MHz), BrukerAvance 400 ($^1$H 400.13 MHz, $^{13}$C 100.61 MHz) and BrukerAvance III 600 ($^1$H 600.25 MHz, $^{13}$C 150.95 MHz) instruments. The spectra are referenced against the NMR-solvent (DMSO-$d_6$: dH = 2.50 ppm) and the chemical shifts δ are reported in ppm. Resonance multiplicity is abbreviated as: s (singlet), d (doublet), t (triplet), q (quartet), m (multiplet) and b (broad). Carbon NMR signals are reported using DEPT135 and $^1$H-$^{13}$C HSQC spectra with (+) for primary/tertiary, (-) for secondary and (q) for quaternary carbons. An Agilent Q-TOF 6540 UHD (ESI-MS) instrument was used for recording mass spectra. UV/Vis absorption spectroscopy was accomplished using a Varian Cary Bio 50 UV/Vis spectrometer in 10 mm quartz cuvettes. IR-spectra were recorded on an Agilent Cary 630 FT-IR spectrometer and the peak positions are reported in wavenumbers (cm$^{-1}$). Analytical HPLC measurements were performed on an Agilent 1220 Infinity LC (column: Phenomenex Luna 3 µM C18(2) 100 Å, 150 x 2.00 mm; flow: 0.3 mL/min at 30 °C; solvent A: MilliQ water with 0.05% vol TFA; solvent B: MeCN). The ratios at the photostationary states (PSSs) were determined by HPLC measurements with a detection wavelength at the isosbestic point. For determination of the thermal half-lives the solutions were irradiated until the photostationary state was reached. Then the solutions were left at 25 °C and the recovery of the absorbance of the *trans*-isomer at $\lambda_{max}$was measured. Consequently, the thermal half-life was calculated by fitting the data with a single exponential function. An Agilent 1260 system (column: Phenomenex Luna 10 µM C18(2) 100 Å, 250 x 21.2 mm; flow: 22.0 mL/min; solvent A: MilliQ water with 0.05% vol TFA; solvent B: MeCN) was used for preparative HPLC purification. Light sources for illumination were: λ = 365 nm (Herolab hand-held lamp UV-6 L, 6 W), λ = 455 nm (OSRAM Oslon SSL 80 LED, 700 mA, 1.12 W), and λ = 530 nm (CREE-XP green, 700 mA, 3.7 W). The power of the light is given based on the specifications supplied by the company when the lamps were purchased. Final compounds for biological testing possess a purity ≥95% determined by HPLC measurements with detection at 220 nm and 254 nm.

**(*E*)-7-((1*H*-pyrazol-4-yl)diazenyl)-5-phenyl-1,3-dihydro-2*H*-benzo[*e*][1,4]diazepin-2-one (3a)**

4-Nitroso-1*H*-pyrazole (**1a,** 71 mg, 0.73 mmol, 1.2 eq.)was added to a solution of amino-benzodiazepine (**2,** 153 mg, 0.61 mmol, 1.0 eq.) in glacial acetic acid (8 mL). Then the mixture was stirred 3 days at room temperature, quenched by adding a saturated aqueous solution of $NaHCO_3$ (30 mL) and diluted with EtOAc (30 mL). After separation of the layers, the aqueous layer was extracted with EtOAc (3x20 mL). The combined organic layers were dried over $Na_2SO_4$, filtered and the solvent was removed under reduced pressure. Purification by preparative HPLC (15% - 45% MeCN in 25min, $t_R$ = 18.8 min) afforded the desired product (30 mg, 15%) as yellow solid. $R_f$ 0.55 ($CH_2Cl_2$/MeOH 9:1); m.p. 133 °C; $^1$H-NMR (600 MHz, DMSO-$d_6$) δ = 13.21 (bs, 1H), 10.86 (s, 1H), 8.22 (bs, 2H), 7.98 (dd, *J* = 8.7, 2.3 Hz, 1H), 7.56 (d, *J* = 2.3 Hz, 1H), 7.55 – 7.51 (m, 3H), 7.48 – 7.43 (m, 2H), 7.39 (d, *J* = 8.8 Hz, 1H), 4.21 (s, 2H); $^{13}$C-NMR (151 MHz, DMSO-$d_6$) δ = 170.0 (q), 169.7 (q), 158.1 (+), 157.9 (+), 146.7 (q), 141.2 (q), 140.8 (q), 138.6 (q), 130.6 (+), 129.4 (+), 128.4 (+), 126.4 (q), 125.0 (+), 124.4 (+), 122.2 (+), 56.9 (−); IR (neat) ν = 3138, 2866, 1707, 1674, 1614, 1487, 1439, 1394, 1346, 1200, 1133, 995, 839, 798, 723 cm$^{-1}$; HRMS (ESI) calculated. for $C_{18}H_{15}N_6O$ $(M+H)^+$ m/z = 331.1302; found 331.1303.

**(*E*)-4-((2-Oxo-5-phenyl-2,3-dihydro-1*H*-benzo[*e*][1,4]diazepin-7-yl)diazenyl)benzenesulfonic acid (9) (3b)**

A 1:1 mixture of tetrabutylammonium 4-nitrosobenzenesulfonate (**1b**) and its corresponding nitro derivative (236 mg, 0.54 mmol, 1.0 eq. of nitroso compound) was added to a solution of amino benzodiazepine (**2,** 68 mg, 0.27 mmol, 1.0 eq.) in acetic acid (2 mL) and $CH_2Cl_2$ (1 mL). After stirring the mixture for 24 h at room temperature the solvent was removed *in vacuo*. Purification by automated flash column chromatography ($CH_2Cl_2$/MeOH, 3% - 25% MeOH) and subsequent preparative HPLC (2% - 65% MeCN in 10 min, $t_R$ = 7.2 min) yielded **3b** (69 mg, 61%) as yellow solid. $R_f$ 0.03 ($CH_2Cl_2$/MeOH 9:1); m.p. 280 °C (decomposition); $^1$H-NMR (600 MHz, DMSO-$d_6$) δ = 11.32 (s, 1H), 8.23 (dd, *J* = 8.8, 2.4 Hz, 1H), 7.81 – 7.78 (m, 2H), 7.77 – 7.73 (m, 3H), 7.68 (t, J = 7.4 Hz, 1H), 7.64 (dd, J = 8.3, 1.3 Hz, 2H), 7.57 (t, J = 7.7 Hz, 2H), 7.52 (d, J = 8.9 Hz, 1H), 4.34 (s, 2H); $^{13}$C-NMR (151 MHz, DMSO-$d_6$) δ = 172.5 (q), 168.9 (q), 151.4 (q), 151.1 (q), 146.5 (q), 143.0 (q), 135.8 (q),132.5 (+), 130.7 (+), 128.8 (+), 128.6 (+), 126.8 (+), 126.2 (+), 124.4 (q), 122.9 (+), 122.2 (+), 54.6 (−); IR (neat) ν = 3489, 3135, 2930, 1715, 1614, 1484, 1435, 1387, 1342, 1230, 1163, 1115, 1029, 1006, 846, 742, 697 cm$^{-1}$; HRMS (ESI) calculated. for $C_{22}H_{16}N_4O_4S$ $(M+H)^+$ m/z = 421.0965; found 421.0964.

**(*E*)-4-((2-Oxo-5-phenyl-2,3-dihydro-1*H*-benzo[*e*][1,4]diazepin-7-yl)diazenyl)benzene-sulfonamide (3c)**

Freshly prepared 4-nitrosobenzenesulfonamide (**3c,** 238 mg, 1.38 mmol, 3.0 eq.) was added to a solution of amino benzodiazepine trifluoroacetate salt (168 mg, 0.46 mmol, 1.0 eq.) in $CH_2Cl_2$ (6 mL) and acetic acid (2 mL). After stirring the mixture for 24 h at 40 °C the solvent was removed *in vacuo*. The residue was purified by automated flash column chromatography ($CH_2Cl_2$/MeOH, 3% - 10% MeOH) yielding **3c** (140 mg, 73%) as orange solid. Material for analytical characterization as well as for biological testing was further purified by preparative HPLC (10% - 75% MeCN in 18 min, $t_R$ = 11.1 min).

$R_f$ 0.14 ($CH_2Cl_2$/MeOH 97:3); m.p. 207 °C; $^1$H-NMR (400 MHz, DMSO-$d_6$) $\delta$ = 11.08 (s, 1H), 8.16 (dd, $J$ = 8.8, 2.3 Hz, 1H), 7.98 (s, 4H), 7.79 (d, $J$ = 2.3 Hz, 1H), 7.58 – 7.46 (m, 8H), 4.27 (s, 2H); $^{13}$C-NMR (101 MHz, DMSO-$d_6$) $\delta$ = 170.1 (q), 169.8 (q), 153.2 (q), 146.2 (q), 146.0 (q) 143.0 (q), 138.0 (q), 131.1 (q), 129.7 (+), 128.5 (+), 127.8 (+), 127.1 (+), 126.1 (q), 124.9 (+), 123.0 (+), 122.6 (+), 56.5 (−); IR (neat) $\nu$ = 3243, 3071, 1700, 1674, 1610, 1484, 1390, 1334, 1200, 1163, 1014, 902, 842, 798, 746, 697 $cm^{-1}$; HRMS (ESI) calculated. for $C_{21}H_{18}N_5O_3S$ $(M+H)^+$ m/z = 420.1133; found 420.1125.

**(*E*)-5-Phenyl-7-(pyridin-2-yldiazenyl)-1,3-dihydro-2*H*-benzo[*e*][1,4]diazepin-2-one (3d) (Glyght)**

2-Nitrosopyridine (**1d,** 108 mg, 1.00 mmol, 2.0 eq.) was added to a solution of amino-benzodiazepine (**2,** 126 mg, 0.50 mmol, 1.0 eq.) in $CH_2Cl_2$ (3 mL) and acetic acid (1 mL). After stirring the mixture for 24 h at room temperature the solvent was removed *in vacuo*. The residue was purified by automated reversed phase flash column chromatography (MeCN/$H_2O$ with 0.05% TFA, 5% - 100% MeCN) and subsequent preparative HPLC (10% - 60% MeCN in 20 min, $t_R$ = 13.4 min) yielding **Glyght** (125 mg, 73%) as yellow solid. $R_f$ 0.63 ($CH_2Cl_2$ + 1% $Et_3N$/MeOH 9:1); m.p. 222 °C; $^1$H-NMR (600 MHz, DMSO-$d_6$) $\delta$ = 10.98 (s, 1H), 8.67 (ddd, $J$ = 4.7, 1.8, 0.8 Hz, 1H), 8.17 (dd, $J$ = 8.7, 2.3 Hz, 1H), 8.00 (ddd, $J$ = 8.1, 7.4, 1.8 Hz, 1H), 7.79 (d, $J$ = 2.2 Hz, 1H), 7.68 (dt, $J$ = 8.0, 1.0 Hz, 1H), 7.56 – 7.51 (m, 4H), 7.48 – 7.44 (m, 3H), 4.25 (s, 2H); $^{13}$C-NMR (151 MHz, DMSO-$d_6$) $\delta$ = 170.1 (q), 169.3 (q), 162.6 (q), 149.4 (+), 146.1 (q), 143.0 (q), 138.9 (+), 138.8 (q), 130.5 (+), 129.3 (+), 128.4 (+), 126.8 (+), 126.7 (q), 125.8 (+), 125.1 (+), 122.4 (+), 113.3 (+), 57.2 (−); IR (neat) $\nu$ = 3105, 3058, 2930, 2881, 1707, 1610, 1487, 1424, 1327, 1245, 1174, 1111, 936, 846, 790, 753, 701 $cm^{-1}$; HRMS (ESI) calculated. for $C_{20}H_{16}N_5O$ $(M+H)^+$ m/z = 342.1349; found 342.1358.

#### *In vitro e*lectrophysiological studies.

##### Cell culture and ion channel expression.

*In vitro* testing of the compound was performed using a line of Chinese hamster ovary (CHO) cells that were transiently transfected with cDNA of different subunits of $GABA_A$ and GlyRs, and $5HT_3$ARs. CHO cells were cultured as previously described[7,8]. Transfection was performed with Lipofectamine 3000 (Life Technology, USA). The following receptor combinations were investigated during this study: α1 zebrafish and α2 zebrafish homomeric GlyRs,α1human and α2 mouse homomeric GlyRs, α1 human/β mouse and α2 mouse/β mouse heteromericGlyRs; heteromeric $GABA_A$Rs formed by human α1/β2/γ2 subunits. Identification of transfected cells was assured by simultaneous transfection of cDNA of green fluorescent protein (GFP); α1 subunit of $GABA_A$ receptor contained GFP as part of the construct. Electrophysiological recordings were performed in the fluorescent cells 24-72 hours after transfection.

##### Electrophysiological recordings in cell lines.

Whole-cellpatch-clamp recordings were held at room temperature (20-25 °C) using an EPC-9 amplifier (HEKA Elektronik, Germany). Cells were continuously superfused with external solution containing (mM): NaCl 140, $CaCl_2$ 2, KCl 2.8, $MgCl_2$ 4, HEPES 20, glucose 10; pH 7.4; 320-330 mOsm. Intracellular solution used for filling recording patch pipettes contained (mM): KCl - 140, $MgCl_2$ - 2, MgATP - 2, BAPTA (tetrapotassium salt) - 2; pH 7.3 at 20 ºC; 290 mOsm. Pipettes were pulled from borosilicate glass capillaries (Harvard Apparatus Ltd, USA) and had resistances of 5-10 MOhm. For the rapid replacement of the solutions, the fast application system was used. Three parallel rectangular tubes (100x100 µm) were positioned 40-50 µm above the recorded cell. The movement of the tubes was controlled by a computer-driven fast exchange system (SF 77A Perfusion Fast-Step, Warner, USA) allowing a 10–90% solution exchange in 3–5 ms, as measured by open electrode controls (1/10 external solution/water). Cells with low input resistance (<150 MOhm) and a rapid run-down (>30% with repetitive application) were excluded from analysis. UV illumination was provided by computer-driven 365 nm LED (Thorlabs), positioned 5 cm above the recorded cell, the power of UV light was reaching 0.6 $mW/mm^2$ at the level of recording chamber, as determined using an optical power meter (Thorlabs). Electrophysiological recordings were performed with PatchMaster (HEKA Electronic, Germany) software.

##### Electrophysiological data analysis and statistics

To obtain the concentration/response curves the amplitude of evoked currents was plotted against different concentrations of agonists and Glyght (Figure 5ghi), and fitted using a non-linear fitting routine of the Origin 7.5 software (OriginLabs, USA) with the Hill equation:

For glycine: $I = I_{max}/(1+(EC_{50} /[A])^{nH})$

For Glyght: $I = I_{max}/(1+([Glyght]/IC_{50})^{nH})$

Where *I* is the normalized current amplitude induced by the agonist at concentration [A], $I_{max}$ is a maximal current induced at given cell, $n_H$ is the Hill coefficient, $EC_{50}$ or $IC_{50}$ are the concentrations at which a half-maximum response was induced, and [Glyght] is the concentration of Glyght used in the experiment. Ionic current recordings were analyzed with Igor Pro 6.02 and Origin 9.0 software. For statistical analysis paired and unpaired t-tests were used. Data are represented as means ± SEM.

#### Drugs

Commercial drugs were obtained from Sigma-Aldrich (France). Stock solutions of Glyght (10 mM) and picrotoxin (50 mM) were prepared using DMSO and then diluted to the final concentration in extracellular solution. Stock solutions of GABA (1 M), glycine (1 M) and 5HT (10 mM) were prepared using MilliQwater. Non-saturating and saturating concentrations of GABA and glycine were chosen accordingly to the agonist $EC_{50}$ value for given receptor. Mean GABA $EC_{50}$ for α1/β2/γ2 $GABA_A$ receptors was determined to be 8 μM (n = 6); mean glycine $EC_{50}$ for α1Z GlyRs was 35 μM (Supplementary figure 5g, n = 6), for α2Z 70 μM (Supplementary figure 5g; n = 6); for α1β – 73 μM (n = 9), α2β – 130 μM (n = 9). Thus, non-saturating concentration of GABA was 5 μM, of glycine – 20 μM for α1, 50 μM for α2, 30 μM for α1β and 100 μM for α2β. Saturating concentration for $GABA_A$ was 300 μM, for α1 and α2 GlyRs – 300 μM and 500 μM of glycine respectively.

#### Electrophysiology in neuromuscular junction (NMJ)

Diaphragm muscle with the attached phrenic nerve was isolated from mouse and mounted to experimental chamber. Preparation was perfused continuously with an aerated physiological saline solution (mM): NaCl - 125; KCl – 2.5; $CaCl_2$ –2; $NaH_2PO_4$ – 1; $MgCl_2$ – 1; glucose – 11. The pH of solution was adjusted to 7.3 at 20 ºC. To prevent muscle contraction following nerve stimulation, the muscles were transversely dissected 1 hour before experiment. Intracellular recording of end-plate potentials (EPPs) was done with sharp glass microelectrodes (4-9 MΩ) using Axoclamp 900A electrophysiological amplifier (Molecular Devices, CA, USA). The motor nerve was stimulated with electrical pulses of supra-threshold amplitude and 0.1 - 0.2 ms duration with Digitimer DS3 stimulator. EPPs were recorded at low frequency stimulation of motor nerve (0.2 Hz). The resting membrane potential (RMP) of the muscle fiber was monitored through the entire experiment; those experiments that showed significant drop of RMP were not analyzed. Electrophysiological signals were digitized at 5 μs intervals stored and processed off-line with a PC. For analysis of EPP amplitude dynamics the first signal in recording was taken as 100 percent.

#### *In vivo* studies

##### Animal housing and photoswitchable behavioral assays

Tupfel-Lon *Danio rerio* embryos were raised in darkness for 6 days post fertilization (dpf) in UV filtered tap water in Petri dishes (daily cleaned and refilled) at 28.5 ºC. Larvae were recorded and video analyzed using the Zebrabox and Zebralab software (ViewPoint Life Sciences). Briefly, 7 dpf larvae were left undisturbed for 40 minutes in 200 µL fresh UV filtered water and in darkness. Continuously, 100 µL were removed and replaced with a double concentrated treatment solution and data and video recording begun. For the first 20 - 40 minutes, larvae were kept in darkness measuring basal activity, named as the resting period (RP). From the $20^{th}$ minute, 2 minutes 365 nm and 455 nm light changes were applied for 3 consecutive cycles, assuring the solutions to transit between their respective *cis* and *trans* photostationary states. Illumination periods at specific wavelengths lasted for 2 minutes.

##### Data analysis and statistics

Zebrafish tracking was performed in real time and data acquisition integrated one- or two-minute intervals using the Zebralab software (ViewPoint Life Science). Data statistical analysis was performed using GraphPad Prism 6 software. Selective illumination was performed with two ordered based [evenly distributed] arrays of 12 light emitting diodes (LEDs) for each wavelength placed 12 cm afar of the multiwell plate. The light intensities, measured with an optical power meter (model Newport 1916-C), were 5.9 $W \cdot m^{-2}$ for 365 nm (UV) and 2.4 $W \cdot m^{-2}$ for 455 nm (Visible-Blue). Larvae activity was measured as the sum of fast swimming durations over one-minute integration per well (fast swimming time). Distance activity was measured as the sum of swimming distances (in millimeters) during burst activities over one-minute integration. Data were analyzed following Two-way ANOVA (p-value 0.05) and are presented as mean ± standard error of the mean (s.e.m.) with the number of larvae (n) indicated in each case.

**UVAR.** UV/Visible activity ratio was extracted from raw activity data as the swimming ratio between the total of six minutes of UV illumination and the total of six minutes of visible illumination for each drug treatment.

### Modeling studies

#### Receptor Structures

The structure of the homopentameric wild-type $\alpha_1$ZGly receptor ($\alpha_1$ZGlyR) used was thecryo-electron microscopy structure solvedin the presence of Gly (PDB entry 3JAE, corresponding to an open state[9]. Missing side chains and hydrogen atoms were added using the psfgen plugin (version 1.6.4[10]) in VMD[11](version 1.9.2). For α2H GlyR we constructed a homology model of the isoform 2B (UniProt code P23416-2) using the $\alpha_{1Z}$GlyR structure as template (sequence identity = 89.2 %). At the ECD/TMD interface, $\alpha 2_H$GlyR differs by only two residues with respect to α1ZGlyR (85-86 VT → IA).

#### Glyght ligand

The initial structures of the Glyght compound (*cis* and *trans* isomers) were created employing the program Avogadro[12] (version 1.1.1). For each isomer, two 1,4-diazepine ring conformations, M and P, were considered, which differ in orientation (below or above the plane, respectively) of C3 and the phenyl substituent of C5[13].For canonical benzodiazepinesthat bind to the classical allosteric site of the $GABA_A$ receptor, the M conformation is the bioactive one[13] (i.e. the one that shows higher affinity for receptor). However, it is not known a priori whether Glyght would exhibit similar conformational preferences, since here we consider binding to a different receptor (GlyR) and to distinct site(s). All four ligand structures (cis/M, cis/P, trans/M and trans/P) were optimized using Density Functional Theory[14] (DFT), with the B3LYP functional[15] and the 6-31++G (d,p) basis set. Calculations were performed with the Gaussian 09 (G09) program package[16]. For the trans isomer, the two conformers differ only by 0.02 kcal/mol and thus their Boltzmann populations are expected to be very similar (50.8% and 49.2% for P and M, respectively). In other words, the two conformers can be present at room temperature. In the case of the cis isomer, the M conformer is significantly more stable (by 1.5 kcal/mol) than the P conformer, and thus it is the predominant conformer (with a Boltzmann population of 92%).

#### Docking calculations

Autodock Vina[17] (version 1.1.2) was employed for ligand-receptor docking. The maximum energy difference between the best and worst binding modes and the exhaustiveness were set to default values (3kcal/mol and 8, respectively). Instead, the maximum number of modes was increase to 20 in order to increase the docking sampling. This protocol was repeated 10 times, starting with different random seeds, so that a total number of 200 binding modes was obtained for each of the four possible conformers of Glyght (cis/M, cis/P, trans/M and trans/P). For the trans isomer, the docking poses obtained for the corresponding M and P conformations were grouped together to carry out the analysis (since the two conformers are almost isoenergetic, see above), resulting in a total of 400 docking poses were considered. For the cis isomer, the 200 docking poses obtained of each conformer were analyzed separately, and only the most populated conformer, cisM, is discussed in the text.

As the location of the putative binding site(s) for Glyght in GlyR is not known, we designed a multilevel binding site screening approach, in the spirit of reference (www.bonvinlab.org/education/HADDOCK-binding-sites/): (1) blind docking using the whole receptor as a search space, (2) information-driven docking focused on the interfacial site between the extracellular and transmembrane domains identified in (1), in order to refine the docking poses and (3) a flexible docking centered in the intersubunit site found in (2) in where the residues K292, T71, T70 (in one subunit) and S289, T70, A68, S66, P291, P201, Q202, L290, R75, R234, F161, Y239, E69 and T71 (in the adjacent subunit) were allowed to move and adapt to the ligand poses.

**Analysis of the docking results**

The initial blind docking results were analyzed in terms ofthe number density of the ligand poses. Previous studies have successfully used this type of analysis to identify ligand binding sites in other ion channels[18,19]. The underlying assumption is that regions of continuous density (or high occupancy) should represent regions of tighter binding. The number density value was compute using the Volmap plugin[20] of VMD[11]. Namely, each Glyght position was replaced with a normalized Gaussian distribution of width equal to 1.5 Å and the Gaussians were additively distributed on a three-dimensional grid of dimensions 0.5 x 0.5 x 0.5 $Å^3$. For the information-driven docking, we analyzed the most populated pose clusters using the quality threshold algorithm implemented in VMD (https://github.com/luisico/clustering) in order to delineate the specific site within the ECD/TMD interface.(3) The flexible docking results were analyzed in terms of the interactions between the ligand and the receptor. On one hand, statistical analysis of protein residues close to the Glyght docking poses was carried out and the percentage contact frequency was calculated considering that a receptor-ligand contact is present if the protein residue is within 5 Å of Glyght. It is assumed that amino acids with high frequencies pinpoint possible binding residues. On the other hand, the representative structure of the most populated cluster(s) was analyzed using the Binana algorithm[21]. The images of the modelling section were generated with either the UCSF Chimera package[22] or the VMD program[11].

### Supplementary Figures

#### Supplementary Figure 1

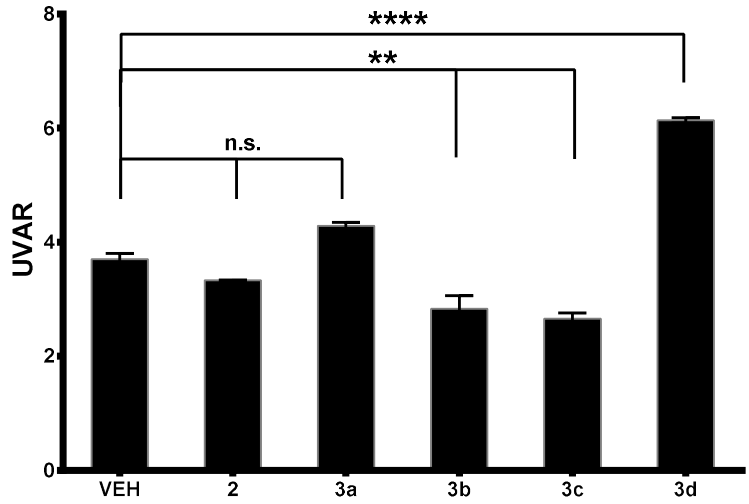


The ratio between activities during UV and visible cycles can be used as a photoswitchable behavior score. The UV/vis activity ratio (UVAR) calculated for each compound can be compared to control UVAR (endogenous photoresponses in vehicle) and provide a simple means to evaluate statistically the significance of the differences. Dunnet's multiple comparison test was conducted (p-value = 0.05).**3b-c** and **3d (Glyght)** were significantly different from vehicle (VEH) with p-values **p < 0.01 and ****p < 0.0001, respectively. In independent experiments, an UVAR = 3.7 ± 0.18 was consistently found in vehicle-treated 7 dpf Tupfel-Lon zebrafish in the described experimental conditions.

#### Supplementary Figure 2

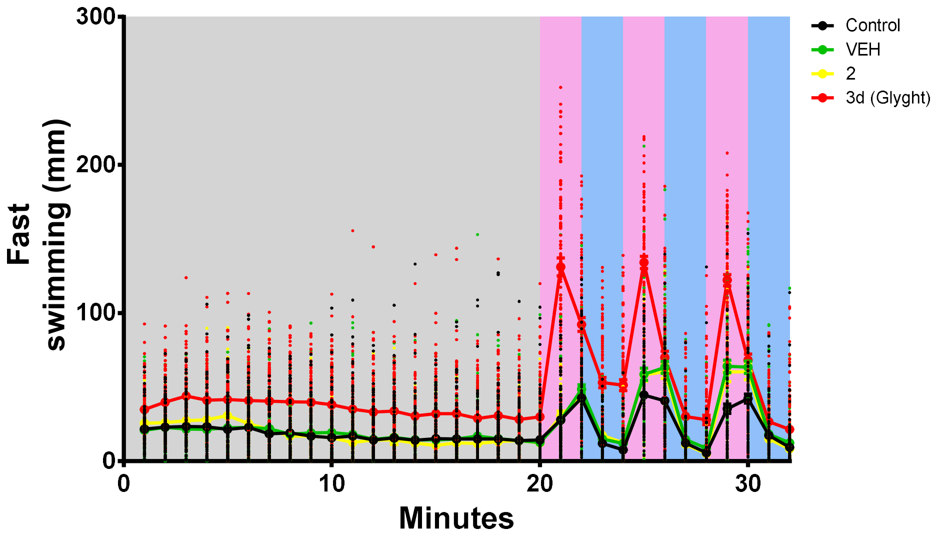


Larvae fast movements activities were analyzed every minute (mm·min$^{-1}$) in five independent experiments –Control (larvae clean water, n = 111), VEH (1%DMSO, n = 95), **2** (n = 46) and **3d** (Glyght) (n = 94),. Experiments record larvae burst depending activity for 20 minutes in Dark condition (RP) followed by 3 consecutives cycles of 2 minutes 365 nm and 455 nm light exposures. Error bars represent S.E.M. for treated larvae over one-minute integration periods.

### Supplementary Figure 3

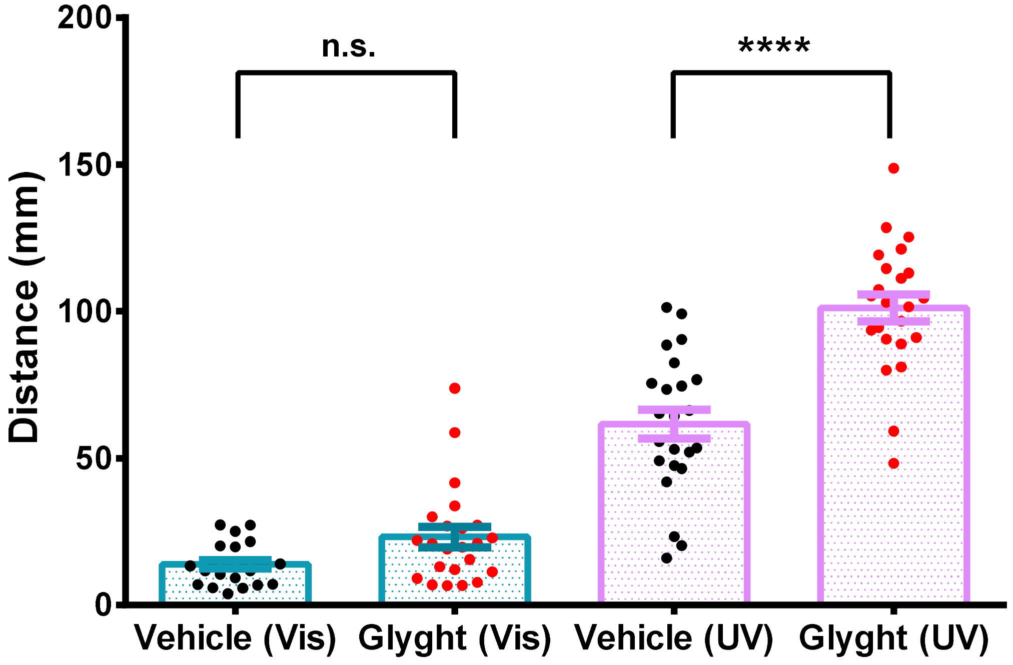


Larvae activity (mm·min$^{-1}$) over all visible and UV light periods was integrated for Vehicle and Glyght treatments (23 larvae per drug treatment). One-way ANOVA withSidak'smultiple comparison test was conducted (p-value = 0.05) Error bars represent S.E.M. Glyght induced a significant increase in larvae activity (p-value < 0.0001) compared to control larvae under UV light, and non-significant (n.s.) differences were found in the same larvae for visible light dependent activities.

### Supplementary Figure 4

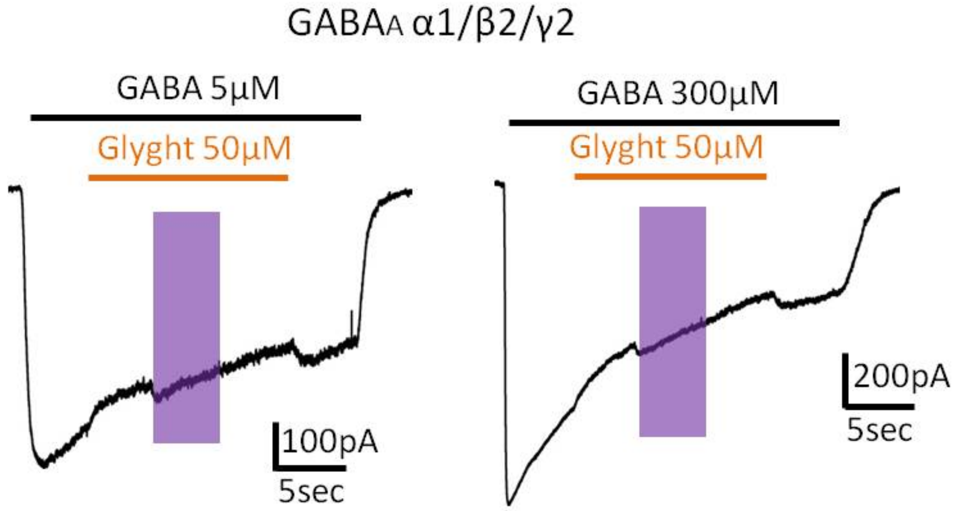


**Glyght has only a minor inhibitory effect on $GABA_A$R-mediated currents.** Left: a representative recording of current induced by application of 5 µM of GABA (black bar) and by mixture of GABA with Glyght 50 µM (orange bar); UV illumination is indicated by violet rectangle. Right: a representative recording of current induced by application of 300 µM of GABA and by mixture of GABA with Glyght 50 µM; $V_{hold}$-30 mV.

### Supplementary Figure 5

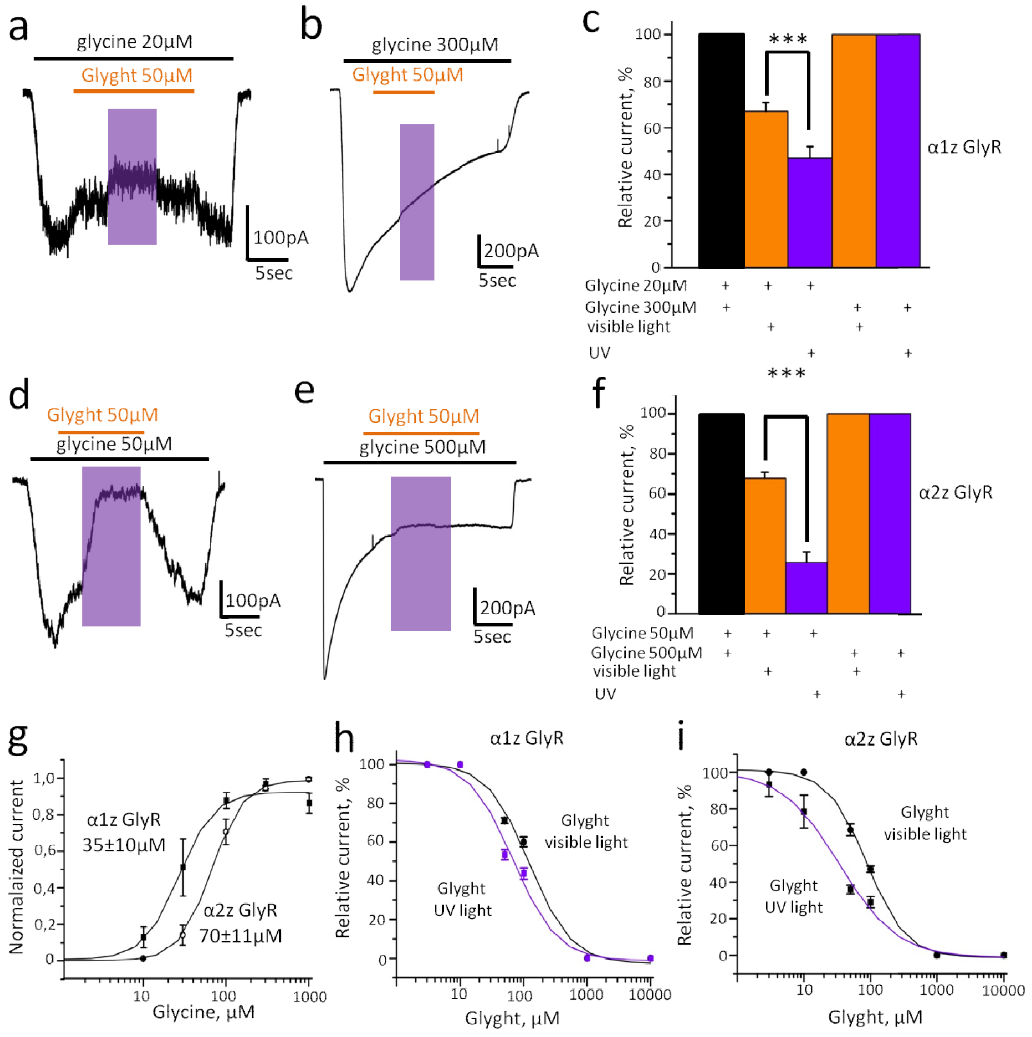


**Glyght UV-dependently inhibits currents mediated by homomeric α1Z and α2Z GlyRs. a,** Representative recording of the α1Z-mediated current induced by application of glycine 20 µM (indicated by black bar) and by mixture of glycine 20 µM and Glyght 50 µM (indicated by orange bar); time of UV illumination indicated by violet rectangle. **b,** Representative recording of the α1Z current induced by 300 µM of glycine and by mixture of glycine 300 µM and Glyght 50 µM. **c,** Relative amplitude of α1Z currents in control (black column; 20 or 300 µM of glycine were applied), at application of the mixture of glycine with Glyght 50 µM under visible light (orange column) and mixture of glycine and Glyght 50 µM under UV light (violet column), $p \leq 0.001$, $n = 8$. **d,** Representative trace of the α2Z current induced by application of glycine 50 µM and by mixture of glycine 50 µM and Glyght 50 µM. **e,** Representative recording of the α2Z current induced by 500 µM of glycine and by mixture of glycine 500 µM and Glyght 50 µM. **f,** Relative amplitude of α2Z currents induced by application of glycine (50 or 500 µM; black column), mixture of glycine with Glyght 50 µM under visible light (orange column) and by mixture of glycine and Glyght 50 µM under UV light (violet column), $p \leq 0.001$, $n = 4$. For all recordings $V_{hold}$-30 mV. **g,**Cumulative dose/response curves for glycine at α1Z (filled squares, n = 6) and α2Z GlyRs (empty circles, n = 6). **h,** Cumulative dose/response curves for Glyght at α1Z GlyRs at visible light (black curve) and upon UV illumination (violet curve); currents were induced by non-saturating concentration of glycine, n = 5. **i,** Cumulative dose/response curves for Glyght at α2Z GlyRs at visible light (black curve) and under UV illumination (violet curve); currents were induced by non-saturating concentration of glycine, n = 4.

### Supplementary Figure 6

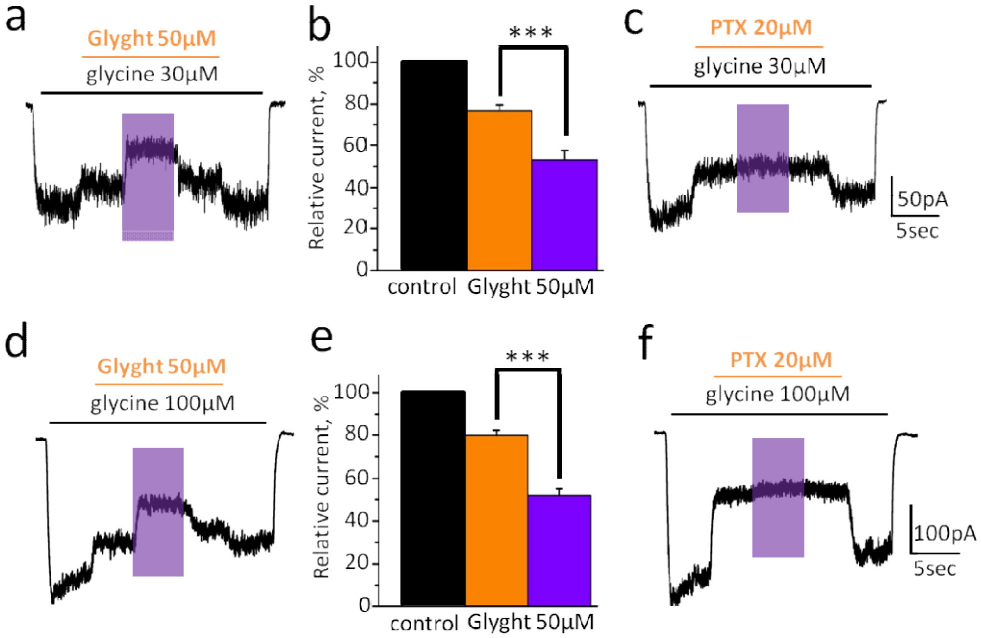


**Glyght inhibited currents mediated by heteromeric α1β and α2β GlyRs in a UV-dependent manner. a,** Representative recording of α1β current induced by 30 µM of glycine and by mixture of glycine 30 µM/ Glyght 50 µM at visible and UV light. **b,** Relative amplitude of α1β currents in control (30 µM of glycine; black column), at application of the mixture of glycine with Glyght 50 µM under visible light (orange column) and mixture of glycine/ Glyght 50 µM under UV light (violet column), n = 7, $p \leq 0.001$. **c,** Representative trace of α1/β current induced by glycine 30 µM and by mixture of glycine with PTX 20 µM, note the absence of the effect of UV light on the current amplitude. **d,** Representative recording of α2β current induced by 100 µM of glycine and by mixture of glycine 100 µM / Glyght 50 µM at visible and UV light. **e,** Relative amplitude of α2β currents in control (100 µM of glycine; black column), at application of the mixture of glycine with Glyght 50 µM under visible light (orange column) and mixture of glycine/ Glyght 50 µM under UV light (violet column), n = 7, $p \leq 0.001$. **f,** Representative trace of α1/β current induced by 100 µM of glycine and by mixture of glycine with 20 µM of PTX illustrating the absence of UV light effect on PTX induced inhibition.

### Supplementary Figure 7

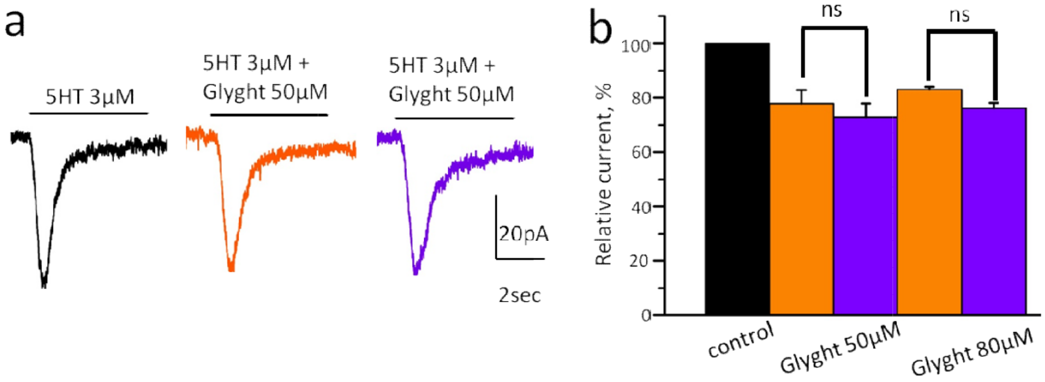


**Glyght does not modulate the amplitude of currents mediated by $5HT_{3A}$ receptors. a,** Representative traces of currents induced by application of 5HT 3 µM (black trace), by mixture of 5HT with Glyght 50 µM at visible light (orange trace) and by mixture of 5HT with Glyght 50 µM at UV light (violet trace). **b,** Cumulative data on relative amplitude of 5HT-induced currents in control (black column), at application of 50 and 80 µM of Glyght at visible light (orange columns) and at application of 50 and 80 µM of Glyght at UV light (violet columns), $p > 0.05$, n = 4.

### Supplementary Figure 8

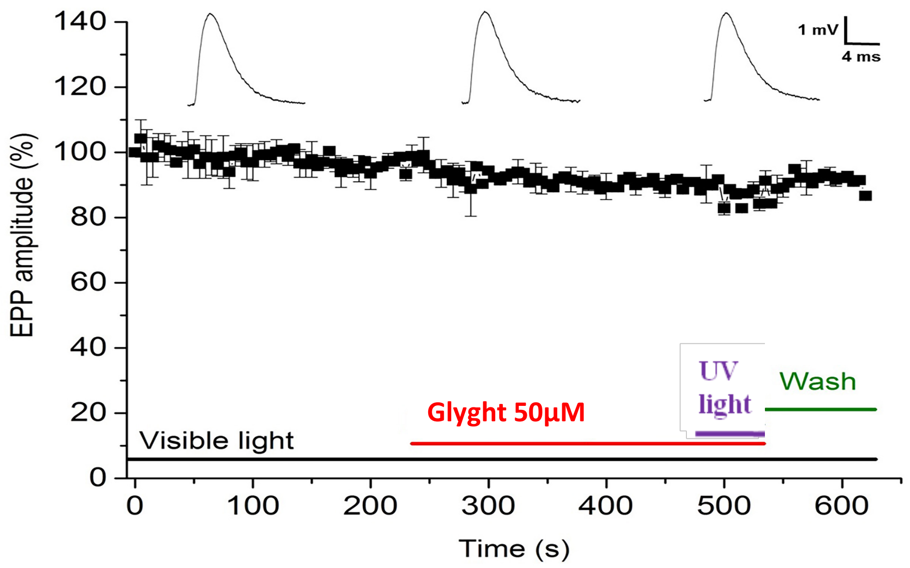


**Testing the influence of Glyght on the amplitude of end-plate potentials at the neuromuscular junction.** The effect of 100 µM of Glyght (shown by red line) on the amplitude of end-plate potentials (EPP) at the mouse neuromuscular junction when applied under the r light (shown by black line) and UV light (365 nm, shown by violet line). Glyght wash-out was done at the end of experiment (shown by green line). At the upper part of the figure are shown examples of native end-plate potential records at different moments of experiment: in control (left), during application of Glyght at visible light (middle) and at UV illumination (right).

### Supplementary Figure 9

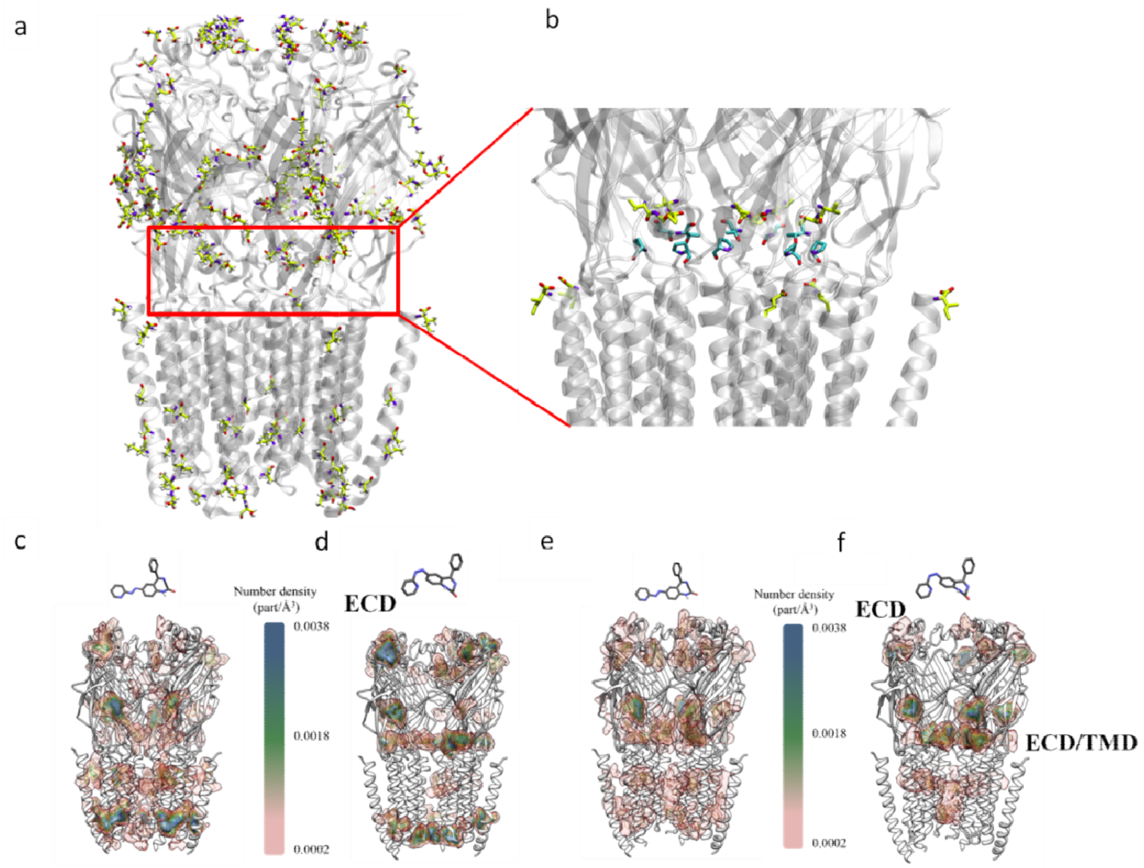


**a, b. Density map of the ligand poses of Glyght obtained in the blind docking with the $\alpha_{1Z}$ GlyR cryo-EM structure (Du et al., 2015), which represents an open state.** Each contour line corresponds to a number density of 0.0006 particles/Å$^3$. **a,** *Trans* isomer (conformers M and P); **b,** *Cis* isomer (conformer M). For the sake of clarity, the front subunit is not shown. **c,d. Density map of the ligand poses of Glyght obtained in the blind docking of the $\alpha_2$ GlyR homology model (open state).** Each contour line corresponds to a number density of 0.0006 particles/Å$^3$. **c,** *Trans* isomer (conformers M and P); **d,** *Cis* isomer (conformer M). For the sake of clarity, the front subunit is not shown. **e,f. ECD/TMD interface involved in the allosteric coupling between ligand binding to the ECD and opening of the ion channel pore in the TMD**[23–27]. Highly conserved amino acid residues throughout pGLICs[27], and residues whose mutation (either in natural variants or site-directed mutagenesis experiments) affect channel activation and conductance[28,29] are shown in ball and sticks colored in white (C atoms), blue (N atoms) and red (O atoms).

### Supplementary Figure 10

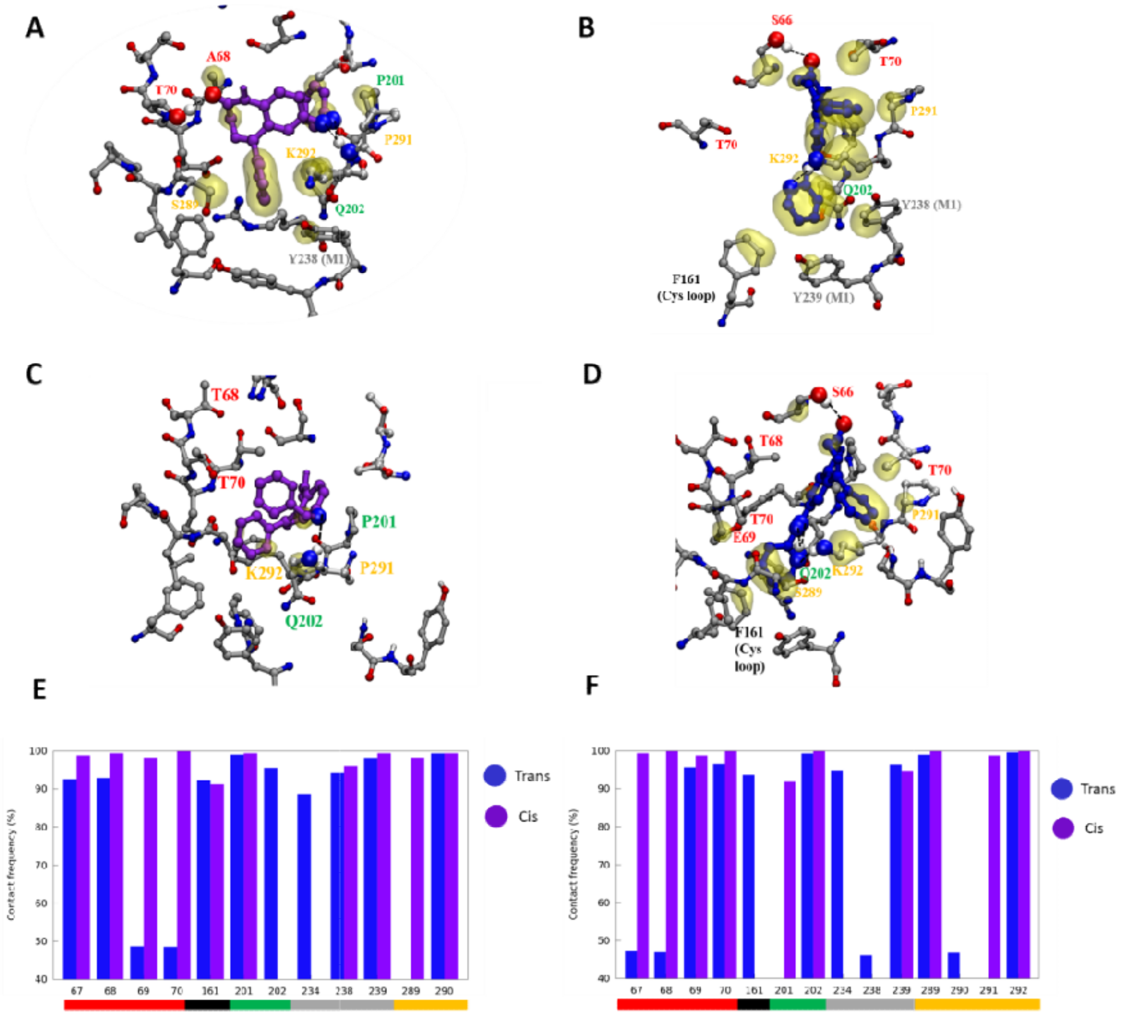


| | | | | | | | |
|---|---|---|---|---|---|---|---|
| Disease related | R287 | K292 | Y295 | V296 | Q242 | | |
| Mutagenesis | A68 | | | | | | |
| Conserved | F161 | P162 | D164 | F203 | R234 | P246 | P291 |

**a, Detailed view of the most representative pose of *cis*-Glyght bound to the ECD/TMD interfacial site obtained in the flexible docking with$\alpha_1$GlyR**. Hydrophobic contacts between *cis*-Glyght and the receptor residues are represented as a yellow surface and the atoms involved in hydrogen bonds between the ligand and Lys292 and Thr70 are represented with more voluminous balls and a dashed line. The residues in direct contact with *cis*-Glyght are indicated. **b, Detailed view of the most representative pose of *trans*-Glyght bound to the ECD/TMD interfacial site obtained in the flexible docking with α1 GlyR**. Hydrophobic contacts between *trans*-Glyght and the receptor residues are represented as a yellow surface and the atoms involved in the hydrogen bonds between the ligand and Lys292 and Ser66 are represented with more voluminous balls and a dashed line. The residues in direct contact with *trans*-Glyght are indicated. **c, Detailed view of the *cis*-Glyght most probable binding site in $\alpha_2$GlyR (67-68IA → 67-68 VT).** Hydrophobic contacts between *cis*-Glyght and the receptor residues are represented as a yellow surface and the atoms involved in the hydrogen bond between the ligand and Lys292 are represented with more voluminous balls and a dashed line. The residues in direct contact with *cis*-Glyght are indicated. **d, Detailed view of the *trans*-Glyght most probable binding site in α2 GlyR.** Hydrophobic contacts between *trans*-Glyght and the receptor residues are represented as a yellow surface and the atoms involved in the hydrogen bonds between the ligand and Lys292, Gln202 and Ser66 are represented with more voluminous balls and a dashed line. The residues in direct contact with *trans*-Glyght are indicated. **e, Percentage of the frequency that a residue is closer than 5Å with Glyghtin α1 GlyR.** In blue the *trans*-Glyght isomer and in violet, *cis*-Glyght. A coloured box below the residues numbers indicates to which loop they belong; red, black, green, grey and yellow correspond to β1-β2, Cys, β8-β9, preM1-M1 and M2-M3 loops, respectively. **f, Percentage of the frequency that a residue is closer than 5Å with Glyghtin α2 GlyR.** In blue the *trans*-Glyght isomer and in violet, *cis*-Glyght. A coloured box below the residue numbers indicates to whichloop they belong; the same colour code as in pancel (e) is used.The highly conserved residues in the pGLICs family and the residues whose mutation affects negatively the channel activity and conductance[9,28,29]are listed in the table below.

### Supplementary Figure 11

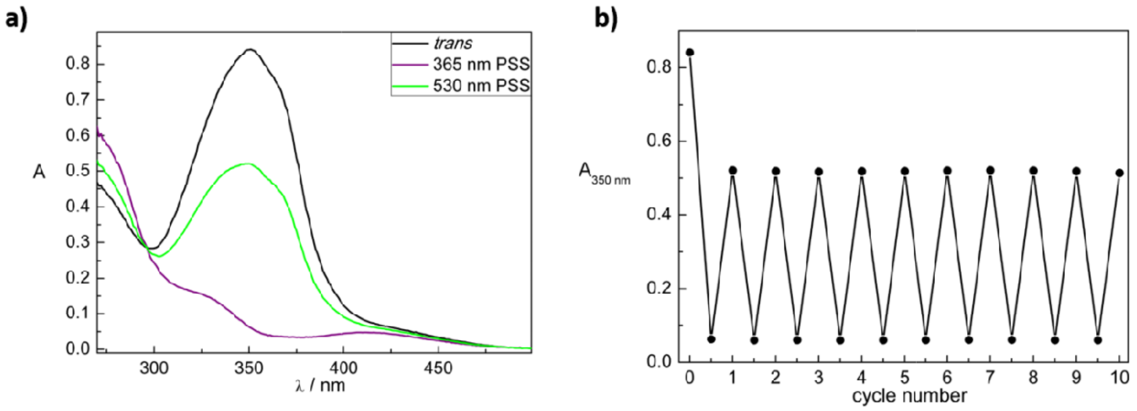


**a,** UV-Vis spectra of **3a** (50 µM in DMSO) from the *trans* isomer (black) at its thermal equilibrium, the PSS at illumination with UV-light of 365 nm (purple) and the PSS at illumination with blue light of 530 nm (green). **b,** Cycle performance of **3a** (50 µM in DMSO). Changes in absorption at 350 nm were measured during alternate illumination with light of 365 nm for 9 s and 530 nm for seven minutes.

### Supplementary Figure 12

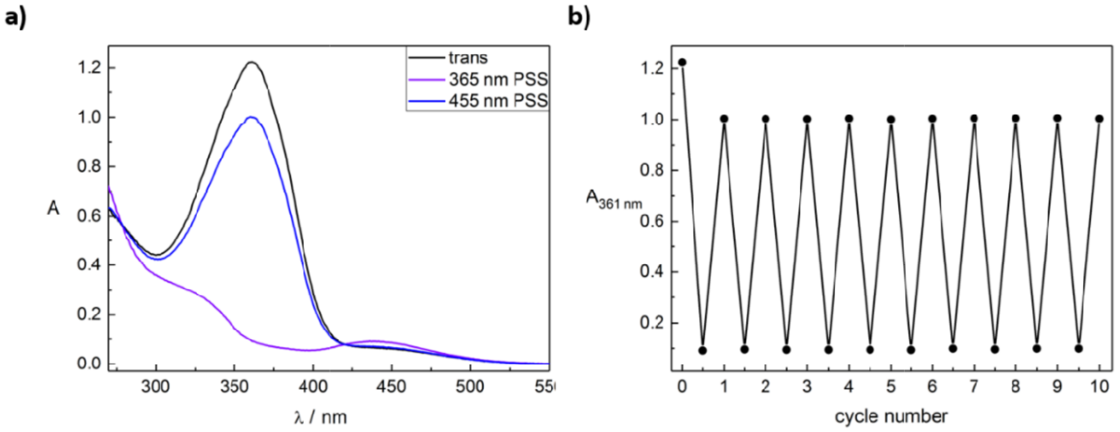


**a,** UV-Vis spectra of azo benzodiazepine **3b** (50 µM in DMSO) from the *trans* isomer (black), the PSS underillumination with UV-light of 365 nm (purple) and the PSS underillumination with blue light of 455 nm (blue). **b,** Cycle performance of azo benzodiazepine **3b** (50 µM in DMSO). Changes in absorption at 361 nm were measured during alternate illumination with light of 365 nm for 15 s and 455 nm for three seconds.

### Supplementary Figure 13

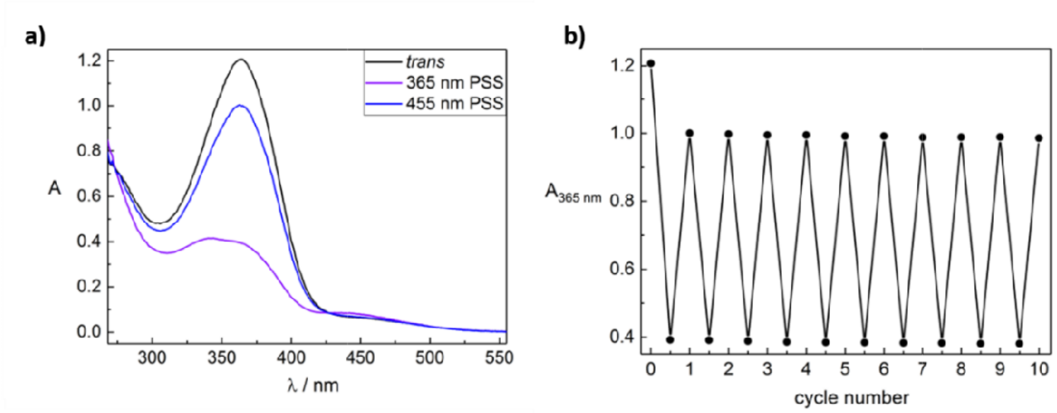


**a,**UV-Vis spectra of **3c** (50 µM in DMSO) from the *trans* isomer (black) at its thermal equilibrium, the PSS underillumination with UV-light of 365 nm (purple) and the PSS underillumination with blue light of 455 nm (blue). **b,** Cycle performance of **3c** (50 µM in DMSO). Changes in absorption at 365 nm were measured during alternate illumination with light of 365 nm and 455 nm.

### Supplementary Figure 14

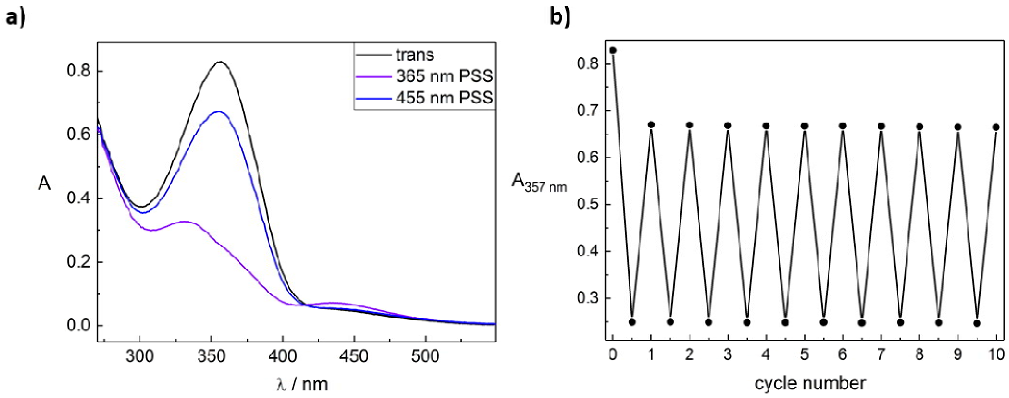


**a,** UV-Vis spectra of **3d** (50 µM in DMSO) from the *trans* isomer (black) at its thermal equilibrium, the PSS under illumination with UV-light of 365 nm (purple) and the PSS under illumination with blue light of 455 nm (blue). **b,** Cycle performance of **3d** (50 µM in DMSO). Changes in absorption at 357 nm were measured during alternate illumination with light of 365 nm for 15 s and 455 nm for three seconds.

### Supplementary Figure 15

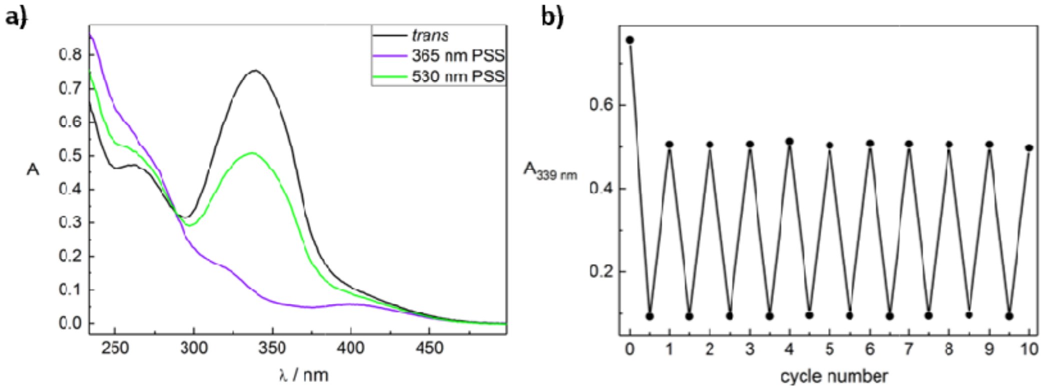


**a,** UV-Vis spectra of **3a**. (50 µM in PBS + 0.1% DMSO) from the *trans* isomer (black), the PSS under illumination with UV-light of 365nm (purple) and the PSS under illumination with blue light of 530nm (green). **b,** Cycle performance (50 µM in phosphate buffer + 0.1% DMSO). Changes in absorption at $\lambda_{max}$ of the *trans* isomer were measured during alternate illumination with light of 365nm and 530 nm.

### Supplementary Figure 16

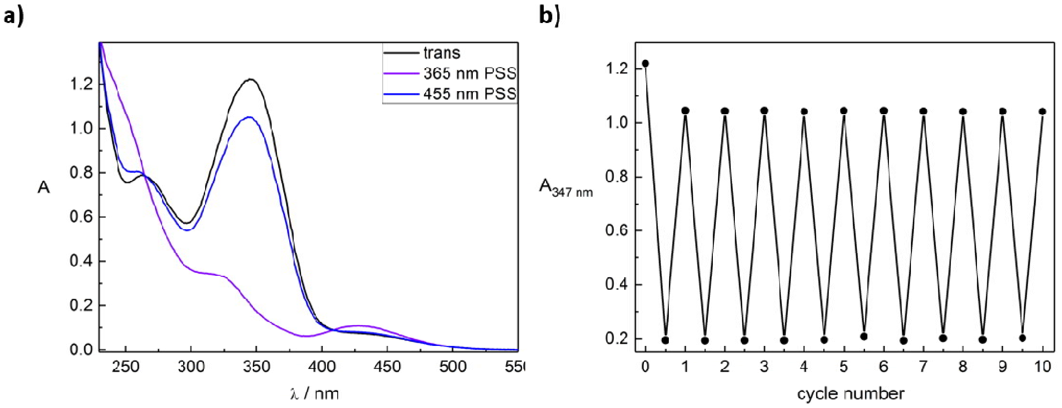


**a,** UV-Vis spectra azo benzodiazepine **3b** (50 µM in PBS + 0.1% DMSO) from the *trans* isomer (black), the PSS under illumination with UV-light of 365nm (purple) and the PSS under illumination with blue light of 455 nm (blue). **b,** Cycle performance of azo benzodiazepine **3b** (50 µM in phosphate buffer + 0.1% DMSO). Changes in absorption at $\lambda_{max}$ of the *trans* isomer were measured during alternate illumination with light of 365 nm and 455 nm.

### Supplementary Figure 17

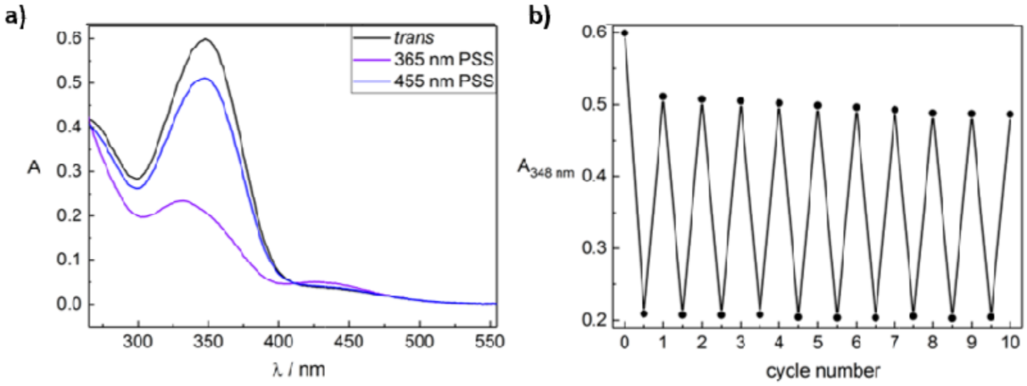


**a,** UV-Vis spectra of azo benzodiazepine **3c** (30 µM in PBS + 3% DMSO) from the *trans* isomer (black), the PSS under illumination with UV-light of 365 nm (purple) and the PSS under illumination with blue light of 455 nm (blue). **b,** Cycle performance (30 µM in phosphate buffer + 3% DMSO). Changes in absorption at $\lambda_{max}$ of the *trans* isomer were measured during alternate illumination with light of 365 nm and 455 nm.

### Supplementary Figure 18

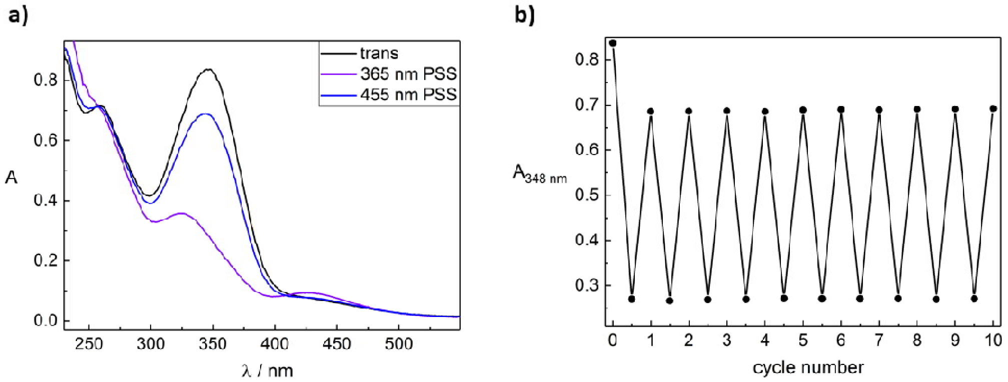


**a,** UV-Vis spectra of azo benzodiazepine **3d** (50 µM in PBS + 0.1% DMSO) from the *trans* isomer (black), the PSS under illumination with UV-light of 365 nm (purple) and the PSS under illumination with blue light of 455 nm (blue). **b,** Cycle performance (50 µM in phosphate buffer + 0.1% DMSO). Changes in absorption at $\lambda_{max}$ of the *trans* isomer were measured during alternate illumination with light of 365 nm and 455 nm.

### Supplementary Figure 19

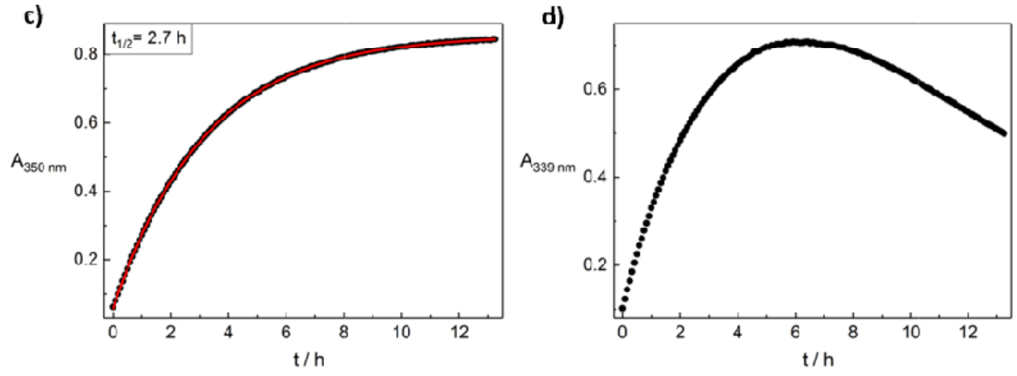


**Compound 3a.** Determination of the thermal half-lives at 25 °C. **Left:** 50 µM in DMSO. **Right:** 50 µM in phosphate buffer + 0.1% DMSO.

### Supplementary Figure 20

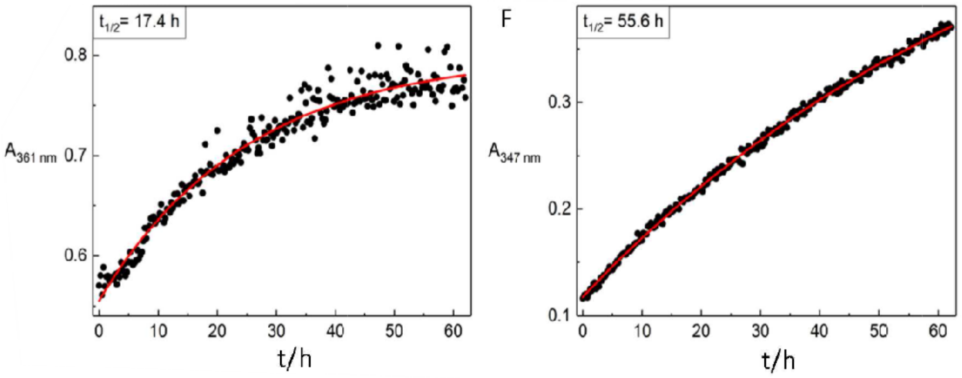


**Compound 3b.** Determination of the thermal half-lives at 25 °C. **Left:** 50 µM in DMSO. **Right:** 50 µM in phosphate buffer + 0.1% DMSO.

### Supplementary Figure 21

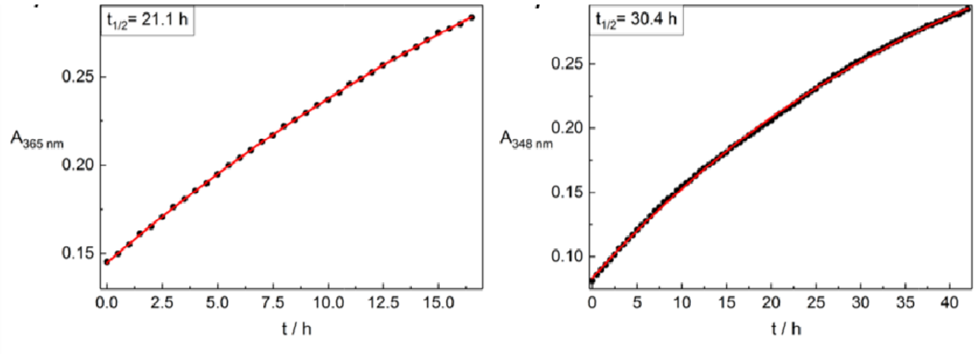


**Compound 3c.** Determination of the thermal half-lives at 25 °C. **Left:** 50 µM in DMSO. **Right:** 50 µM in phosphate buffer + 0.1% DMSO.

### Supplementary Figure 22

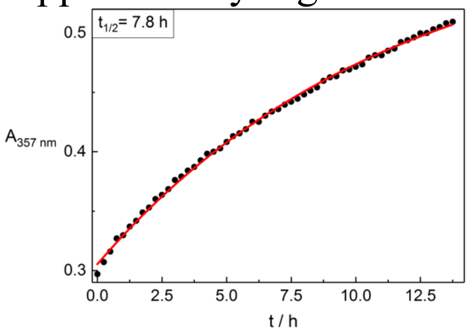


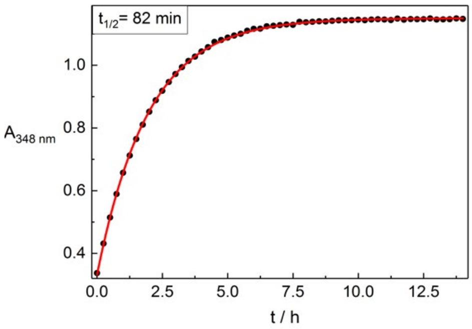


**Compound 3d.** Determination of the thermal half-lives at 25 °C. **Left:** 50 µM in DMSO. **Right:** 50 µM in phosphate buffer + 0.1% DMSO.

### Supplementary Table 1

Photochemical properties of substituted azobenzene benzodiazepine derivatives **3 a-d**.

| Entry | Cpd | Conc. [µM] | solvent | $\lambda_{max}$*trans* [nm] | $\lambda_{max}$*cis* [nm] | $t_{1/2}$(25 °C) [h] | PSS[*,a]*cis* [%] | PSS[*,b]*trans* [%] |
|---|---|---|---|---|---|---|---|---|
| **1** | **3a** | 50 | DMSO | 350 | none | 2.7 h | 88 | 71 |
| **2** | **3a** | 50 | PBS + 0.1% DMSO | 339 | none | - [c] | n.d. | n.d. |
| **3** | **3b** | 50 | DMSO | 361 | 440 | 17.4 | 88 | 82 |
| **4** | **3b** | 50 | PBS + 0.1% DMSO | 347 | 429 | 55.6 | n.d. | n.d. |
| **5** | **3c** | 50 | DMSO | 365 | 442 | 21.1 | 88 | 82 |
| **6** | **3c** | 50 | PBS + 0.1% DMSO | 348 | 427 | 30.4 | n.d. | n.d. |
| **7** | **3d** | 50 | DMSO | 357 | 435 | 7.8 | 80 | 84 |
| **8** | **3d** | 50 | PBS + 0.1% DMSO | 348 | 425 | 1.4 | n.d. | n.d. |

n.d.: not determined; [*]determined by HPLC measurements; [a] PSS at photoconversion from the *trans* to the *cis* isomer; [b] PSS at photoconversion from the *cis* to the *trans* isomer. [c] could not be determined due to decomposition or precipitation after 5 h in PBS solution.

### Analytical HPLC traces for Purity Determination

Compound **3a**
Detection at 220 nm: $t_R$ = 10.8 min; 98%

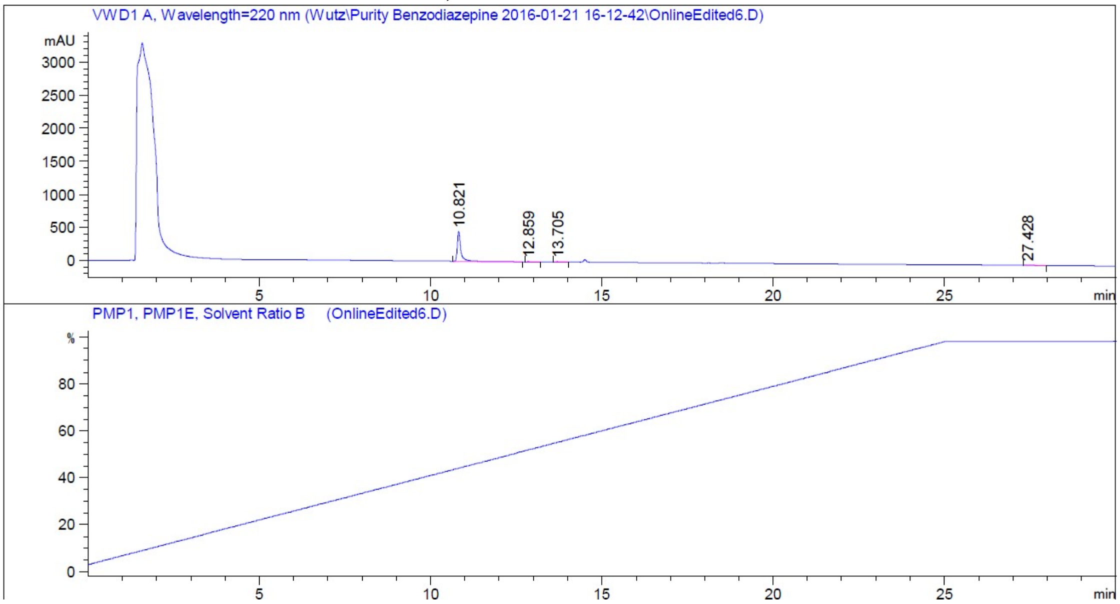


Detection at 254 nm: $t_R$ = 10.8 min; 100%

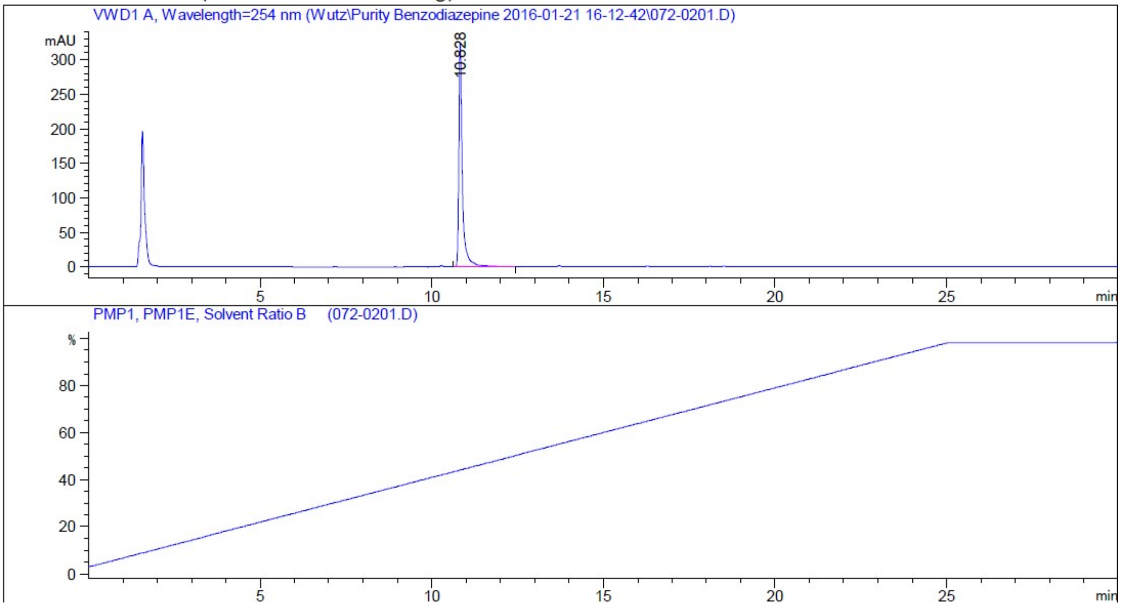

Compound **3b**

Detection at 220 nm: $t_R$ = 9.8 min; 98%

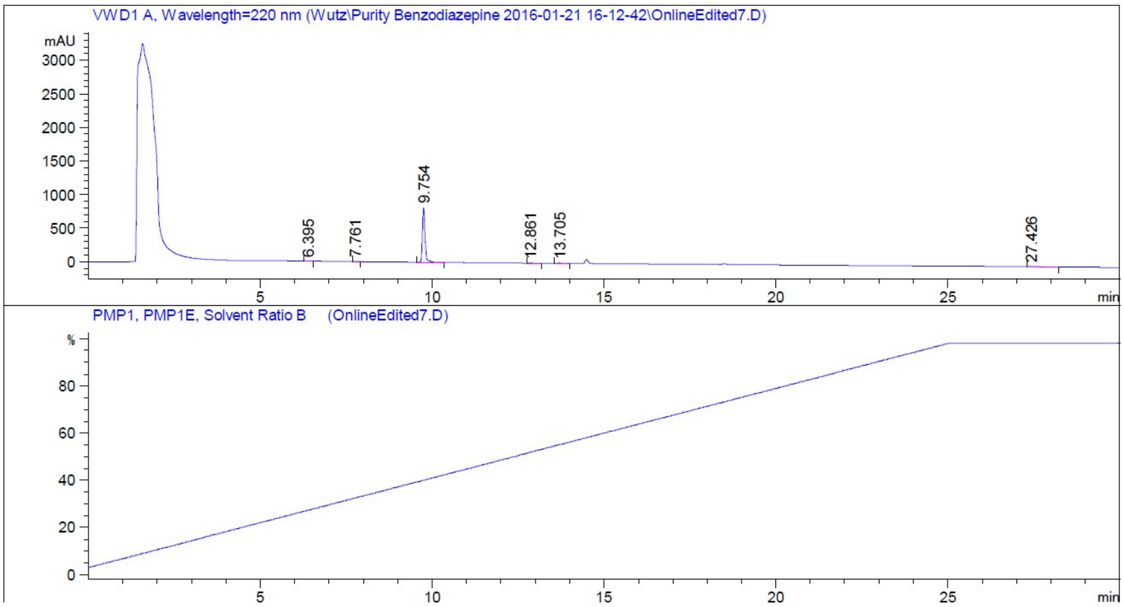


Detection at 254 nm: $t_R$ = 9.8 min; 98%

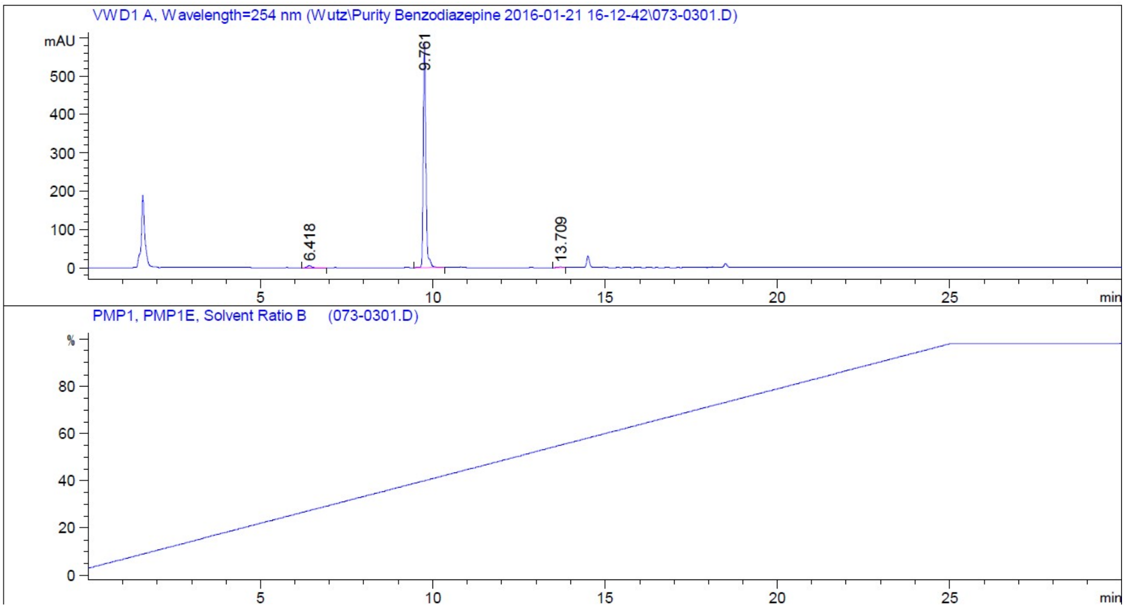

Compound **3c**

Detection at 220 nm: $t_R$ = 13.0 min; 98%

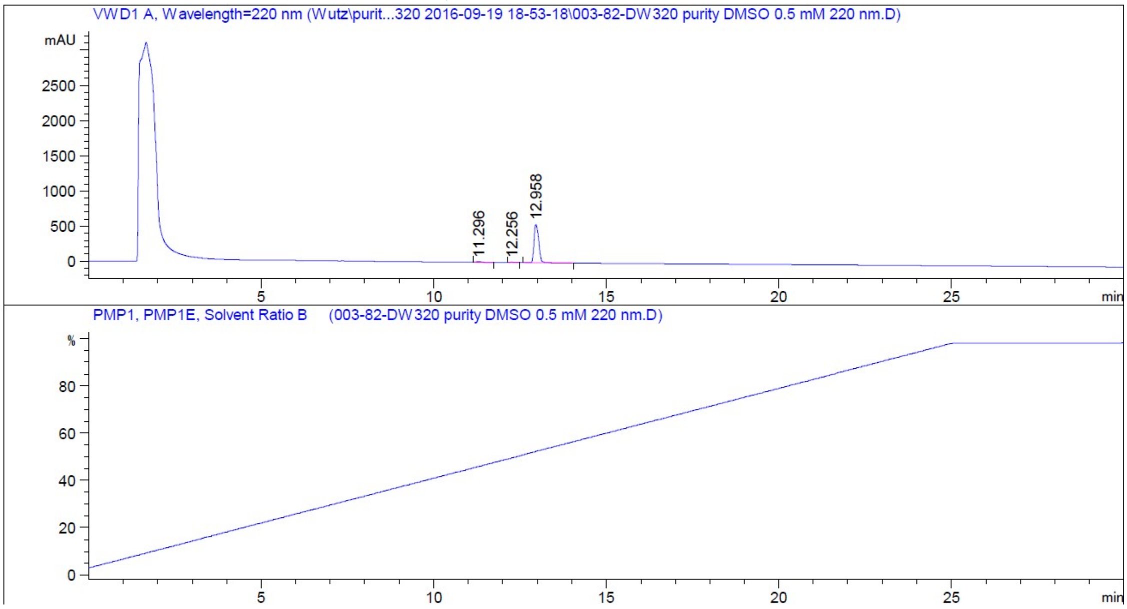


Detection at 254 nm: $t_R$ = 13.0 min; 96%

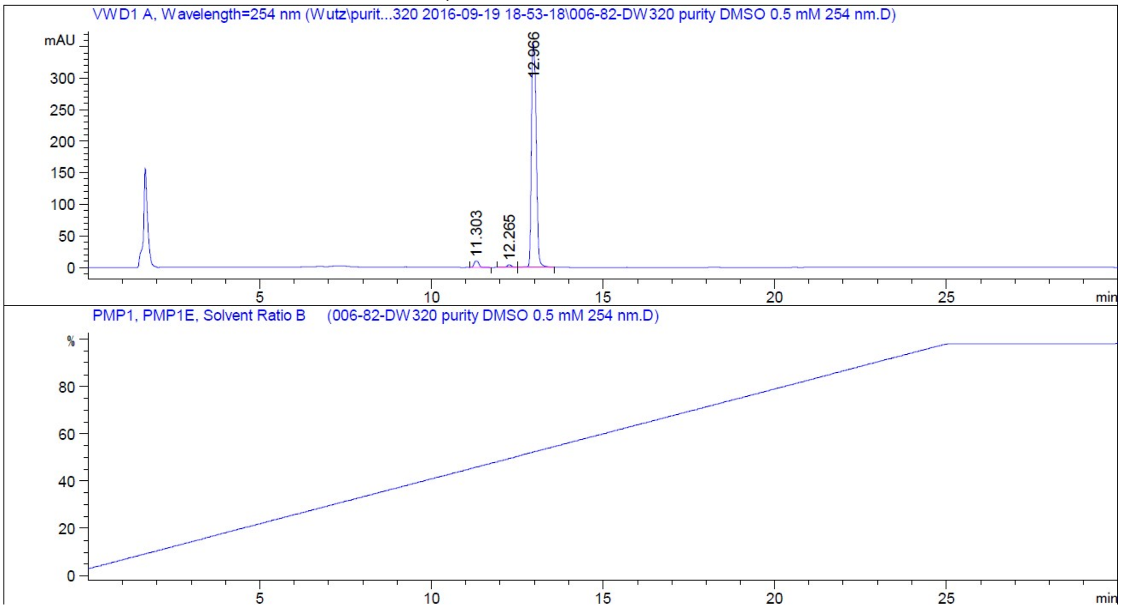

Compound **3d**

Detection at 220 nm: $t_R$ = 13.1 min; 99%

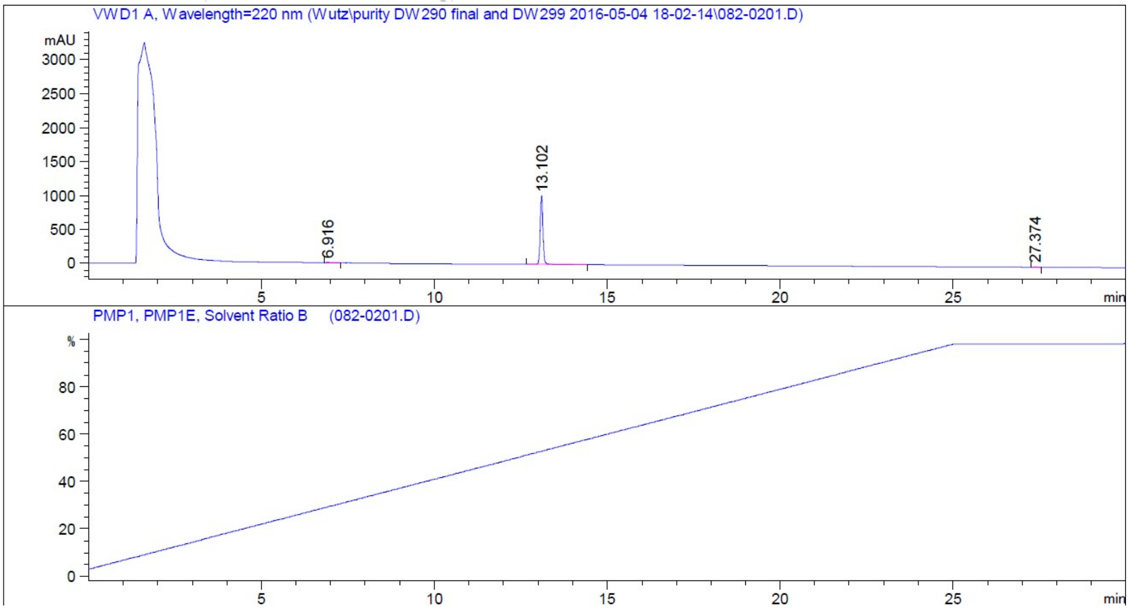


Detection at 254 nm: $t_R$ = 13.1 min; 95%

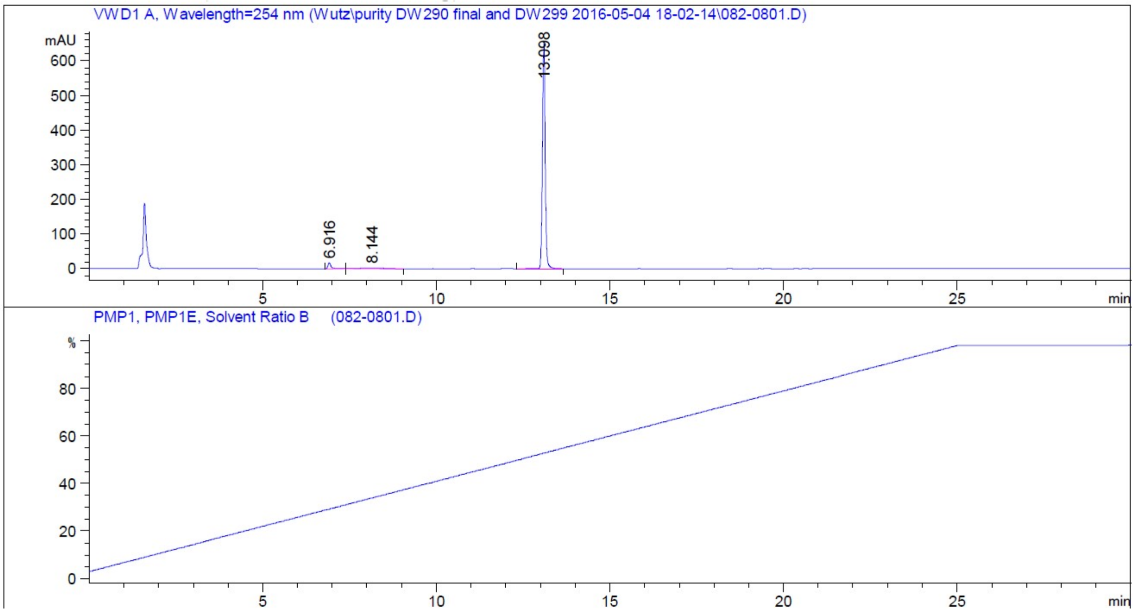

### $^{1}$H- and $^{13}$C-NMR spectra

Compound **3a**

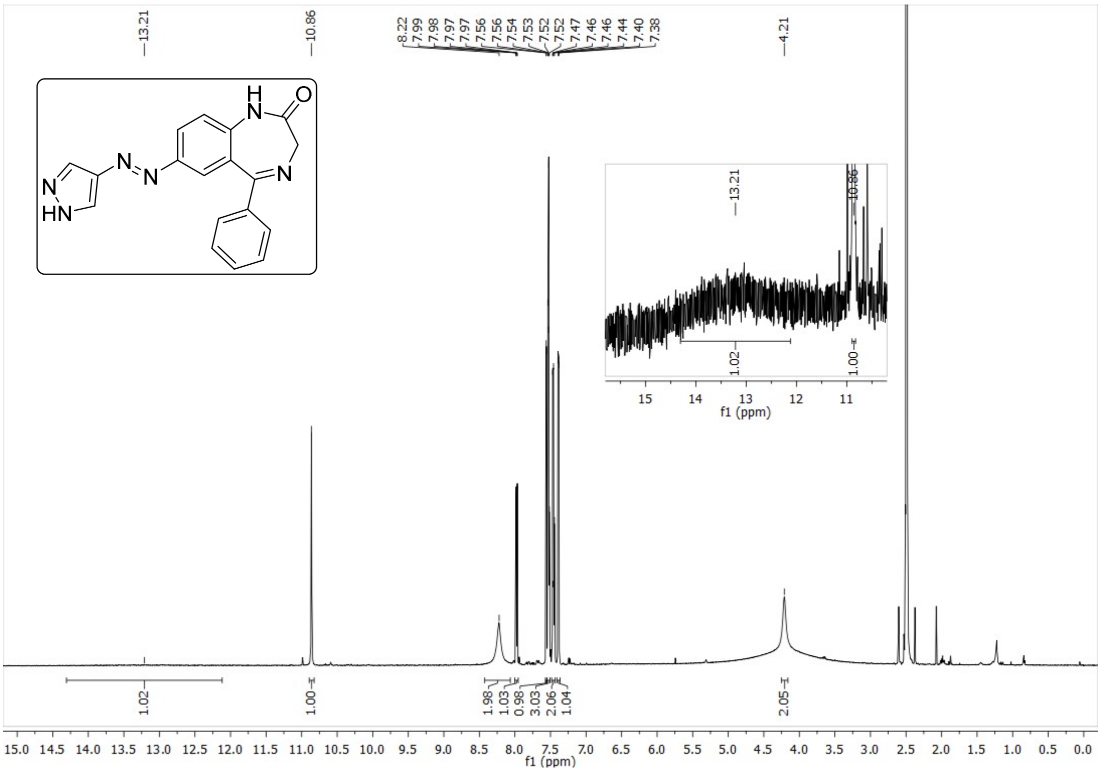

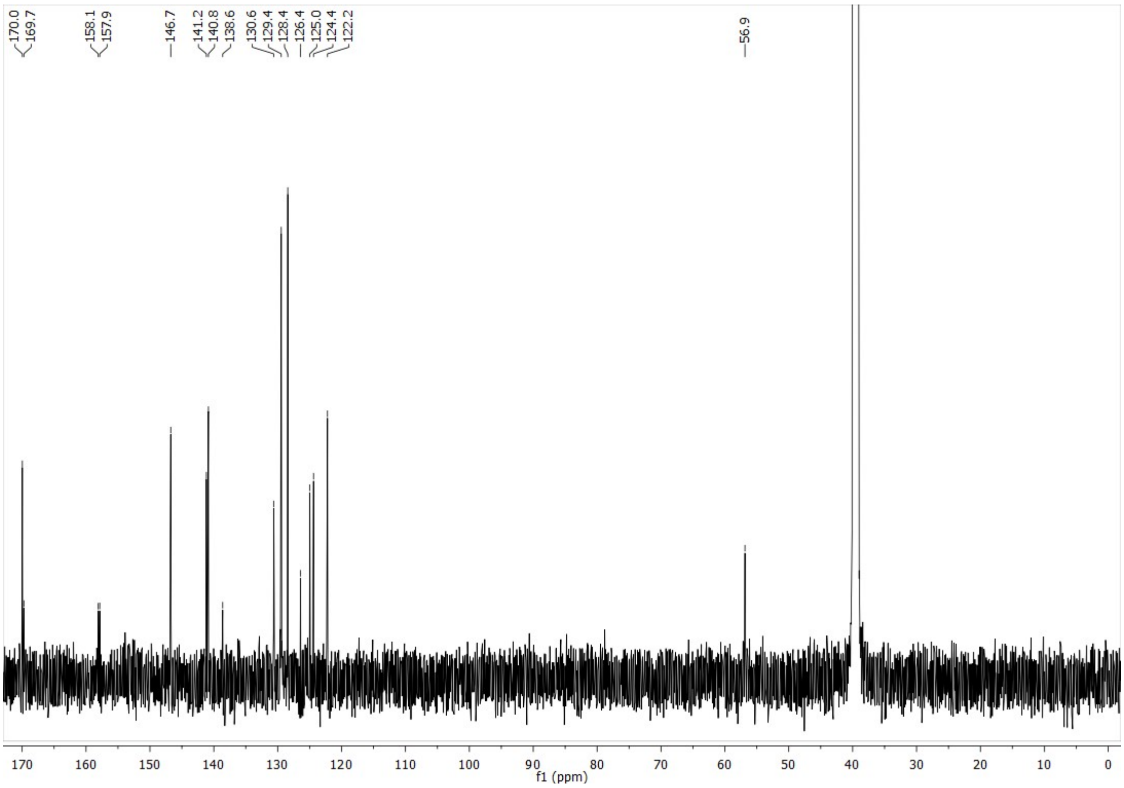

Compound **3b**

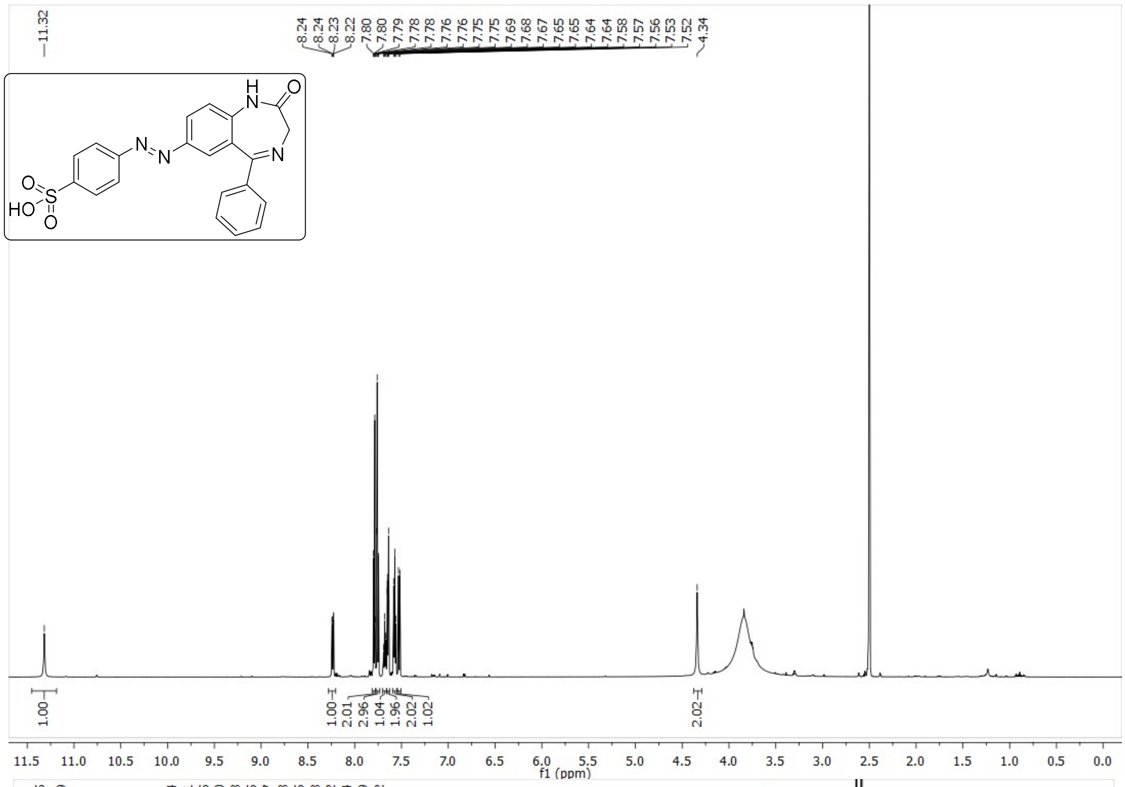
11.32
8.24
8.24
8.23
8.22
7.80
7.80
7.79
7.78
7.78
7.76
7.76
7.75
7.75
7.69
7.68
7.67
7.65
7.65
7.64
7.64
7.58
7.57
7.56
7.53
7.52
4.34
1.00
1.00
2.01
2.96
1.04
1.96
2.02
1.02
2.02
f1 (ppm)

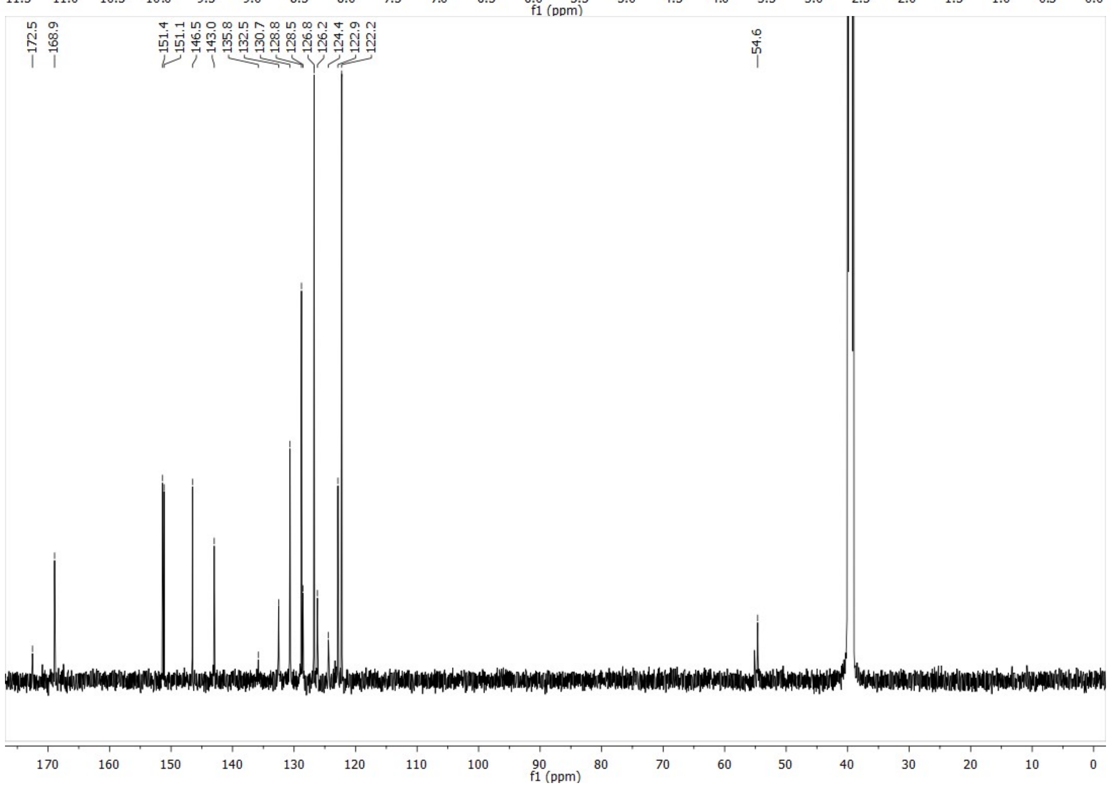
172.5
168.9
151.4
151.1
146.5
143.0
135.8
132.5
130.7
128.8
128.5
126.8
126.2
124.4
122.9
122.2
54.6
f1 (ppm)

Compound **3c**

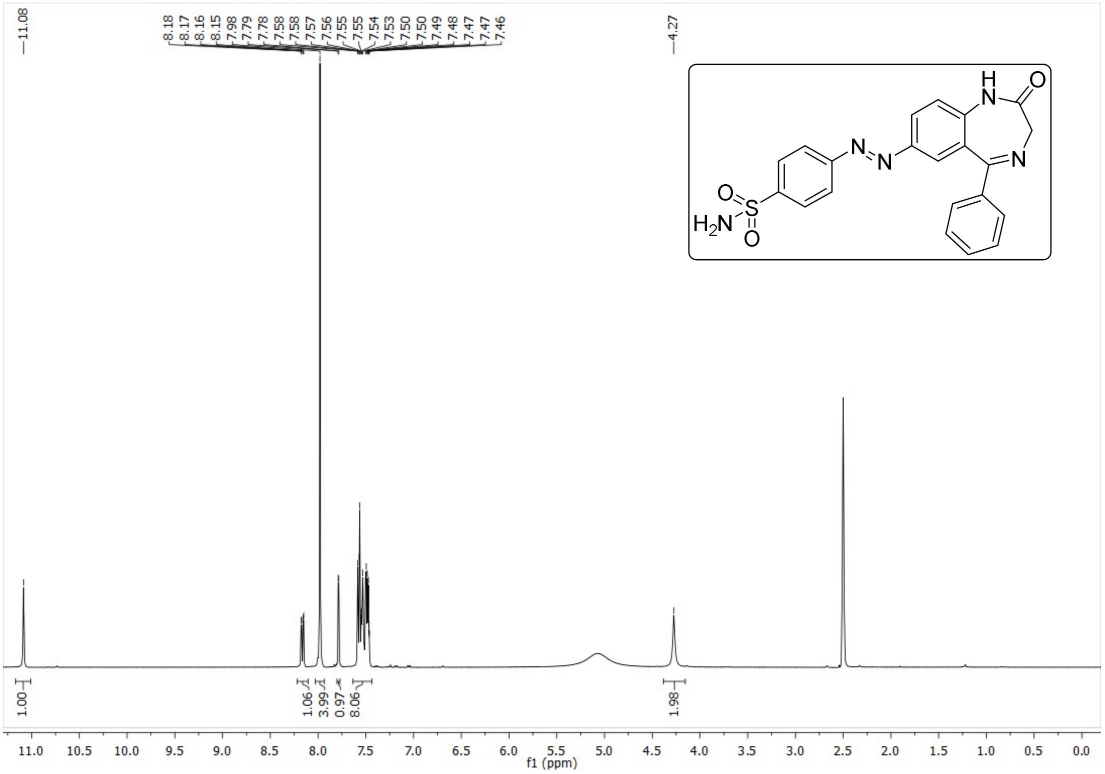
11.08
8.18
8.17
8.16
8.15
7.98
7.79
7.78
7.58
7.58
7.57
7.56
7.55
7.55
7.54
7.53
7.50
7.50
7.49
7.48
7.47
7.47
7.46
4.27
1.00
1.06
3.99
0.97
8.06
1.98
f1 (ppm)
H
N
O
N
N
N
O
S
$H_2N$
O

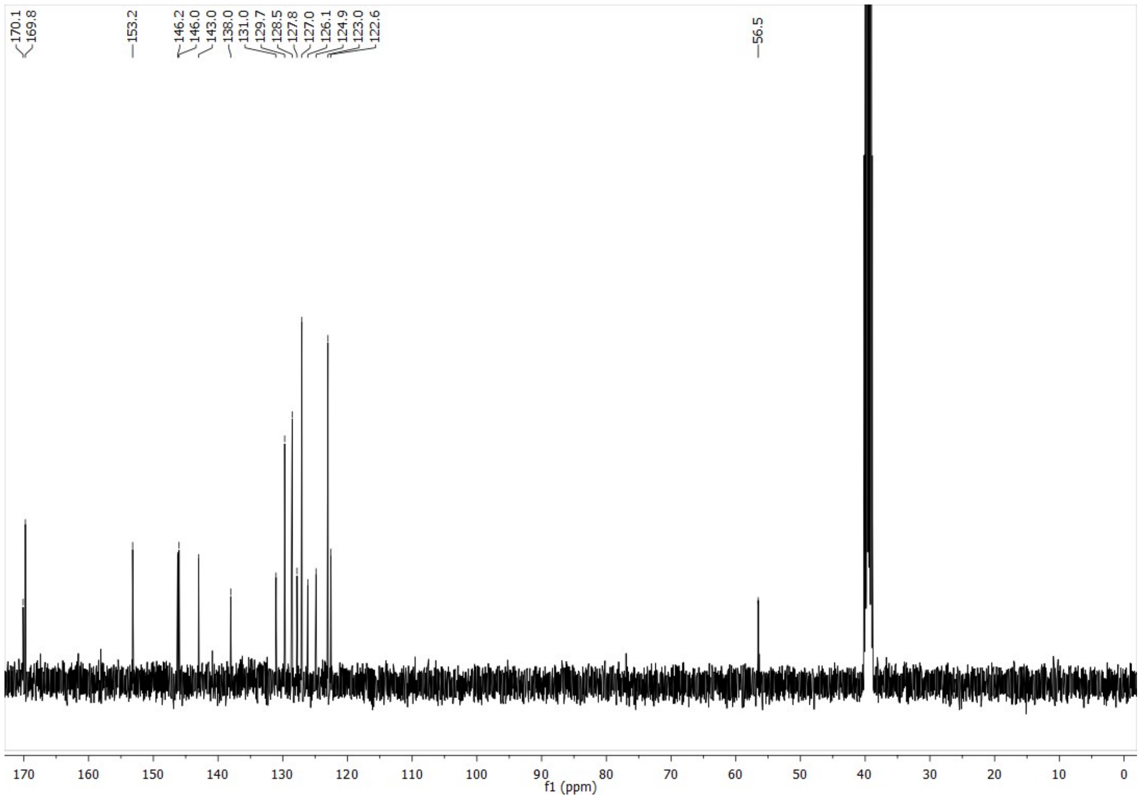
170.1
169.8
153.2
146.2
146.0
143.0
138.0
131.0
129.7
128.5
127.8
127.0
126.1
124.9
123.0
122.6
56.5
f1 (ppm)

Compound **3d**

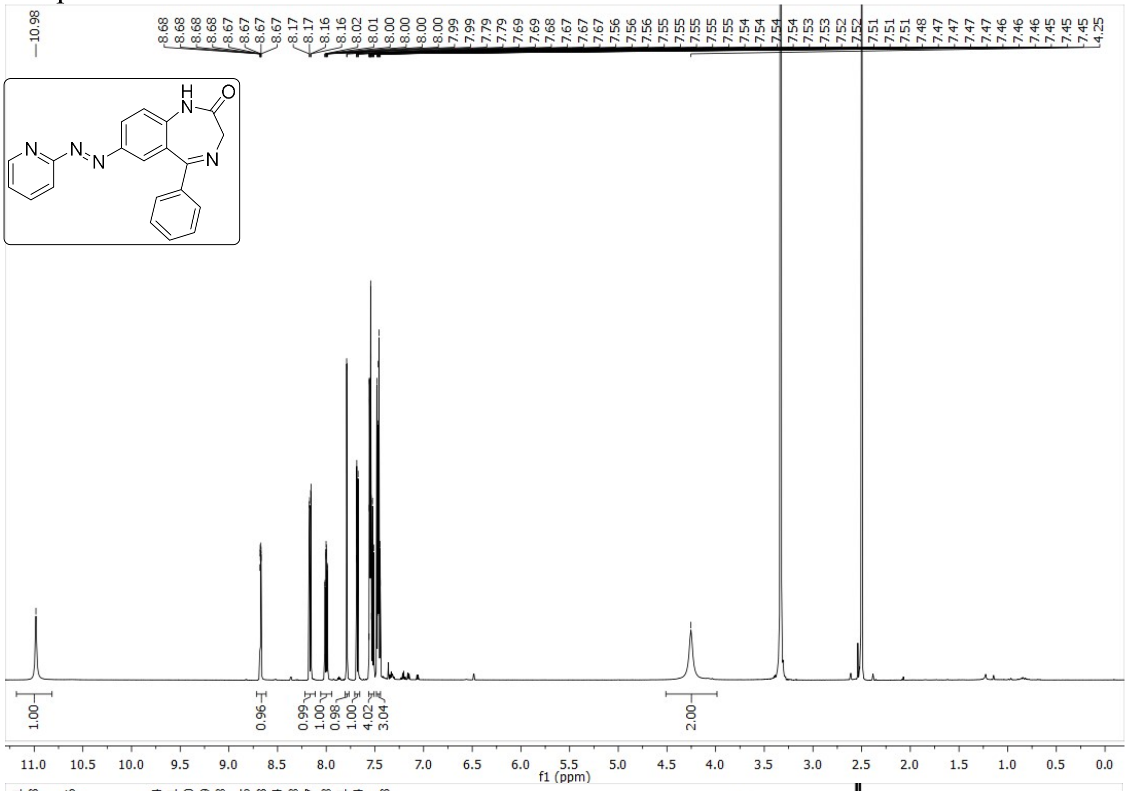
10.98
8.68 8.68 8.68 8.68 8.67 8.67 8.67 8.67 8.17 8.17 8.16 8.16 8.02 8.01 8.00 8.00 8.00 8.00 7.99 7.99 7.79 7.79 7.69 7.69 7.68 7.67 7.67 7.67 7.56 7.56 7.56 7.55 7.55 7.55 7.55 7.55 7.54 7.54 7.54 7.54 7.54 7.53 7.53 7.52 7.52 7.51 7.51 7.51 7.48 7.47 7.47 7.47 7.46 7.46 7.45 7.45 7.45 4.25
H
N
O
N
N
N
N
1.00
0.96
0.99
1.00
0.98
1.00
4.02
3.04
2.00
11.0 10.5 10.0 9.5 9.0 8.5 8.0 7.5 7.0 6.5 6.0 5.5 5.0 4.5 4.0 3.5 3.0 2.5 2.0 1.5 1.0 0.5 0.0
f1 (ppm)

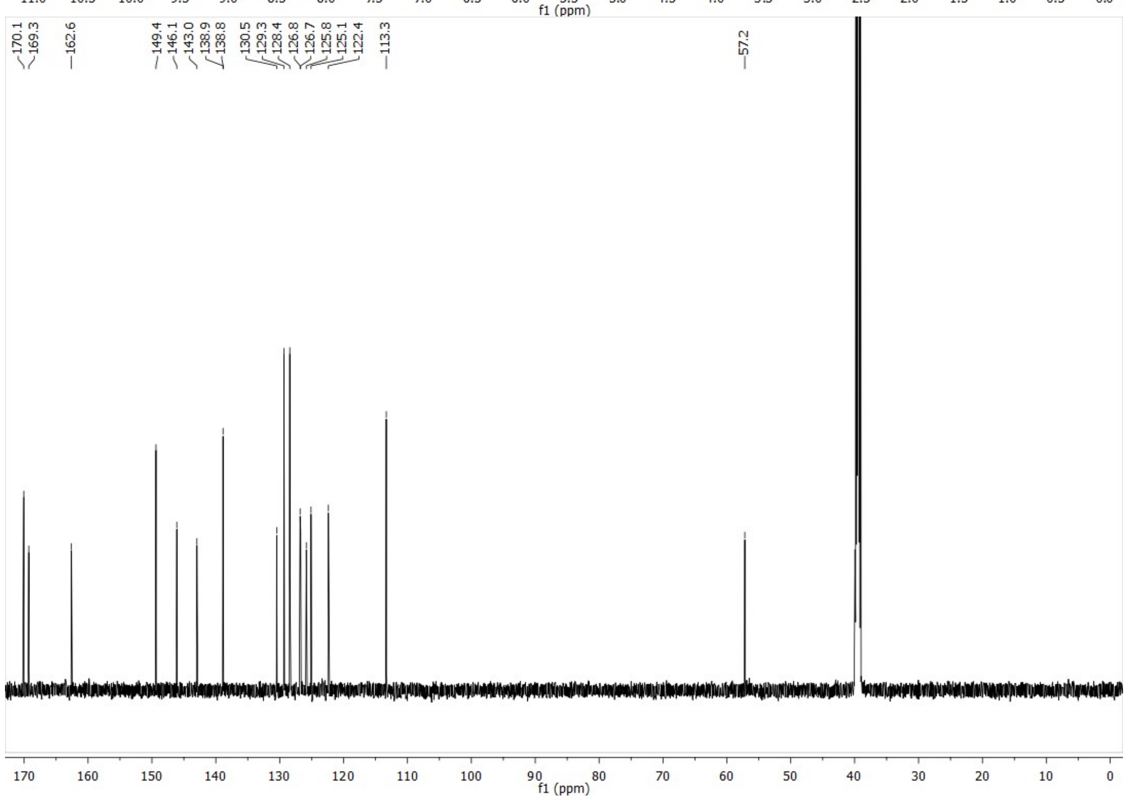
170.1 169.3 162.6 149.4 146.1 143.0 138.9 138.8 130.5 129.3 128.4 126.8 126.7 125.8 125.1 122.4 113.3 57.2
170 160 150 140 130 120 110 100 90 80 70 60 50 40 30 20 10 0
f1 (ppm)

1. Guandalini, L. *et al.* Design, synthesis and preliminary biological evaluation of new hydroxamate histone deacetylase inhibitors as potential antileukemic agents. *Bioorganic Med. Chem. Lett.* **18,** 5071–5074 (2008).
2. Severino, B. *et al.* Synthesis and pharmacological evaluation of peptide-mimetic protease-activated receptor-1 antagonists containing novel heterocyclic scaffolds. *Bioorganic Med. Chem.* **16,** 6009–6020 (2008).
3. Birkofer, L. & Franz, M. C-Nitrosierungen über Silyl-Derivate. *Chem. Ber.* (1971). doi:10.1002/cber.19711041009
4. Priewisch, B. & Rück-Braun, K. Efficient preparation of nitrosoarenes for the synthesis of azobenzenes. *J. Org. Chem.* (2005). doi:10.1021/jo048544x
5. Runtsch, L. S. *et al.* Azobenzene-based inhibitors of human carbonic anhydrase II. *Beilstein J. Org. Chem.* (2015). doi:10.3762/bjoc.11.127
6. Taylor, E. C., Tseng, C. P. & Rampal, J. B. Conversion of a Primary Amino Group into a Nitroso Group. Synthesis of Nitroso-Substituted Heterocycles. *J. Org. Chem.* (1982). doi:10.1021/jo00342a035
7. Mukhtarov, M. *et al.* Calibration and functional analysis of three genetically encoded Cl−/pH sensors. *Front. Mol. Neurosci.* **6,** 1–12 (2013).
8. Maleeva, G., Buldakova, S. & Bregestovski, P. Selective potentiation of alpha 1 glycine receptors by ginkgolic acid. *Front. Mol. Neurosci.* **8,** 1–11 (2015).
9. Du, J., Lü, W., Wu, S., Cheng, Y. & Gouaux, E. Glycine receptor mechanism elucidated by electron cryo-microscopy. *Nature* **526,** 224–9 (2015).
10. Justin Gullingsrud, Jan Saam, J. P. psfgen User's Guide. (2006).
11. Humphrey Dalke Schulten, W. A. K. Visual Molecular Dyanmics. *Journal of Molecular Graphics* **14,** 33–38 (1996).
12. Hanwell, M. D. *et al.* Avogadro: An advanced semantic chemical editor, visualization, and analysis platform. *J. Cheminform.* (2012). doi:10.1186/1758-2946-4-17
13. Richter, L. *et al.* Diazepam-bound GABAA receptor models identify new benzodiazepine binding-site ligands. *Nat. Chem. Biol.* **8,** 455–464 (2012).
14. Rajagopal, A. K. & Callaway, J. Inhomogeneous electron gas. *Phys. Rev. B* (1973). doi:10.1103/PhysRevB.7.1912
15. Stephens, P. J., Devlin, F. J., Chabalowski, C. F. & Frisch, M. J. Ab Initio calculation of vibrational absorption and circular dichroism spectra using density functional force fields. *J. Phys. Chem.* **98,** 11623–11627 (1994).
16. Frisch, M. J. *et al.* Gaussian 09, Revision D.01. *Gaussian Inc.* 1–20 (2013). doi:10.1159/000348293
17. Trott, O. & Olson, A. Autodock vina: improving the speed and accuracy of docking. *J. Comput. Chem.* **31,** 455–461 (2010).
18. Raju, S. G., Barber, A. F., LeBard, D. N., Klein, M. L. & Carnevale, V. Exploring Volatile General Anesthetic Binding to a Closed Membrane-Bound Bacterial Voltage-Gated Sodium Channel via Computation. *PLoS Comput. Biol.* **9,** 1–10 (2013).
19. Bregestovski, P. D. & Maleeva, G. V. Photopharmacology: A Brief Review Using the Control of Potassium Channels as an Example. *Neurosci. Behav. Physiol.* **49,** 184–191 (2019).
20. Cohen, J., Arkhipov, A., Braun, R. & Schulten, K. Imaging the migration pathways for O2, CO, NO, and Xe inside myoglobin. *Biophys. J.* **91,** 1844–1857 (2006).
21. Durrant, J. D. & McCammon, J. A. BINANA: A novel algorithm for ligand-binding characterization. *J. Mol. Graph. Model.* **29,** 888–893 (2011).
